# A genomic and structural bioinformatic pipeline identifies candidate type VI secretion antibacterial effector-immunity pairs

**DOI:** 10.1101/2023.03.26.534264

**Authors:** Alexander Martin Geller, David Zlotkin, Maor Shalom, Noam Blum, Asaf Levy

## Abstract

Type VI secretion systems (T6SS) are common bacterial contractile injection systems that inject toxic “effector” proteins into neighboring cells. We bioinformatically investigated T6SS core proteins in 11,832 genomes of Gram negative bacteria. Comparison of T6SS core proteins that are covalently attached to toxic T6SS effector proteins (T6Es) versus those that are not revealed differences in phylogenetic distribution, physical properties, and genomic position. Using the data generated from our bioinformatic analysis, we developed a new genomic- and Alphafold2-based pipeline for discovery of putative T6Es. We experimentally validated the toxic and immunity activities of four putative antibacterial T6SS effector proteins and four cognate immunity genes from diverse species, respectively. We used Foldseek to predict possible mechanisms of action of the putative T6Es, which was much more effective than sequence-based methods. Evidence of the possible mechanisms of action of the putative T6Es was explored through fluorescence microscopy, where we observed cell wall-targeting, DNA degradation, and cell filamentation. This study shows how combining genomic data mining with new structure-based bioinformatic tools can facilitate identification of novel antibacterial toxins.

## Introduction

Microbes utilize various antagonistic mechanisms to infect hosts and to kill competing microbes. The type VI secretion system (T6SS) is a membrane-bound, contact-dependent secretion system that injects toxic T6SS effector proteins (T6Es) into neighboring bacteria and into eukaryotic cells ^1–4^. Structurally, the T6SS is a contractile injection system, which shoots a spear-like structure towards neighboring cells. The tip and shaft are secreted from the attacking cell ^5^. The tip is made up of a protein called PAAR (Proline-alanine-alanine-arginine), and a trimer of VgrG (Valine-glycine repeat protein G), while the shaft is made up of a column of hollow ring structures, formed from stacks of hexamers of the protein Hcp (Hemolysin-coregulated protein) ^6^.

T6Es are oftentimes proteins that non-covalently interact with Hcp, VgrG, or PAAR, and are thereby delivered upon T6SS contraction and penetration into target cells. These T6Es are sometimes called “cargo” effectors ^7^. However, sometimes the Hcp, VgrG, and PAAR proteins (which here we call T6SS “core proteins”) contain an N-terminal core domain, e.g. a VgrG domain, and a C-terminal toxin domain, e.g. an enzymatic toxin domain. We call core proteins with C-terminal domains “evolved” cores (e.g. “evolved” PAAR). “Evolved” cores are also referred to as “specialized effectors” ^7,8^ (Supplementary Figure 1). An example of an “evolved” core from *Vibrio cholerae* encodes for an “evolved” VgrG with a C-terminal actin crosslinking domain ^9^. Other examples of “evolved” core effectors include VgrG-3 of *V. cholerae*, which has a C-terminal peptidoglycan degrading activity ^10^, VgrG2B of *Pseudomonas aeruginosa*, which has a C-terminal metallopeptidase activity ^11^, Hcp-ET1, which has a C-terminal nuclease domain ^12^, and Tse6 of *P. aeruginosa* has an N-terminal PAAR with a C-terminal toxin ^13,14^.

T6SS and its T6Es are not only important to microbial ecology via their key role in niche colonization and pathogen-host interaction ^15–18^ but also to application as potential new antimicrobials, to medicine and to agriculture^19^. For example, study of T6Es highlights novel bacterial antimicrobial targets. A study of T6E Tse8 showed that inhibition of transamidosome activity has antibacterial activity ^20^. This knowledge can lead to development of new antimicrobials that inhibit a novel molecular target, the transamidosome, to treat infectious diseases. In addition, antifungal effectors were shown to target pathogenic *Candida* species^3,21^, a critical priority human pathogen. Therefore, it is valuable scientifically and clinically to investigate and discover diverse T6Es. Despite this importance, a survey from 2018 screened Proteobacterial genomes that contain T6SS genes and found that only 42% had effectors of known activity, which suggests a large portion of effectors remain undiscovered or their mechanisms of action are yet unknown ^22^.

Generally, T6Es have been discovered in specific model organisms by looking at genetic linkage, i.e. functionally screening genes that are encoded near the T6SS core genes for toxic activity ^23^. Another method to discover T6Es are experimental screens ^23^, but this method has limited scalability, as it requires growth and mutation of the species of interest. An alternative method to identify T6Es is to use bioinformatic approaches to mine the wealth of prokaryotic genomes currently available publicly. Past studies that have used computational approaches to search for new effector genes have relied on algorithms that use 3D protein modeling homology to known enzymatic activities as parameters. The Mougous group has successfully identified a peptidoglycan amidase T6E superfamily using an algorithm that filtered hits for potential amidase activity by protein modeling ^24^, and subsequently used a similar algorithm to search for putative lipase and lysozyme activity, respectively ^25,26^. Another bioinformatic approach used to discover new T6Es was a sequence homology based algorithm, which identified conserved amino acid sequence motifs that are present in a variety of T6SS effectors ^27,28^. A machine learning approach was taken to identify putative T6Es, but the authors did not validate their computational predictions ^29^. Finally, comparative genomics of T6SS-encoding versus non-encoding genomes has also been used as a powerful tool to identify novel T6Es ^30^.

Many T6E-immunity gene pairs remain hidden in microbial genomes, and new methods are required to efficiently discover these genes. In this study, we performed a broad scale bioinformatic analysis on 11,832 T6SS-encoding genomes, with a special focus on “evolved” Hcp, VgrG, and PAAR properties. We explore the characteristics of “evolved” cores in our large genome dataset, and we also use this information to develop a new method to predict and validate novel putative T6E-immunity gene pairs. Our method incorporates Alphafold2, the revolutionary protein folding neural network that has the ability to predict protein folding from primary sequence ^31^. We show how the multimer model of Alphafold2 can be used to predict toxin-antitoxin binding, and how Foldseek, a structure-based search algorithm can help identify possible mechanisms of action of putative T6Es that have little to no sequence-based annotations ^32–35^. Overall, we experimentally identify four putative T6E-immunity protein pairs, and develop models of the modes of action of the new toxins using imaging.

## Results

### Bioinformatic analysis of “evolved” T6SS cores

Toxic T6E proteins are non-covalently or covalently attached to T6SS core proteins to be secreted from the attacking cell into neighboring cells^36^. In order to understand the characteristics of these T6SS cores, we systematically mined and compared “evolved” and “unevolved” T6SS cores in 11,832 bacterial genomes.

Using our dataset of T6SS cores, we asked what proportion of cores are “evolved”, i.e. have a C-terminal domain extension (Materials and Methods). While it is common for PAAR and VgrG to be “evolved” (38.48% and 26.40% have C-terminal domains, respectively), Hcp only quite rarely has “evolved” versions, with only 2.07% of Hcp having a C-terminal domain (Supplementary Figure 2A). We speculate that it may be physically challenging to fit functional toxic effectors into the small (diameter of the central channel is ∼40 Angstroms)^37^, donut-shaped Hcp hexamer for proper packaging, assembly, and delivery. In order to explore whether size constraint may play a role in “evolved” cores, we plotted the sizes of the C-terminal extensions of Hcp, VgrG, and PAAR (Supplementary Figure 2B). Hcp has the smallest range of C-terminal domains (between 71 and 528 amino acids, median size 253 amino acids). The vast majority of VgrG C-termini are also less than 500 amino acids (median size 135 amino acids), yet we observed some outliers of larger size, up to a maximum of 3,663 amino acids. The C-terminal domains of PAAR has a bimodal size distribution, with one group of effector domains of less than 500 amino acids, and another large group of 1200-1400 amino acids. This latter group of large effector domains are virtually all annotated as RHS domains, which are large domains that are found in Nature to have a variety of C-terminal toxins ^38–40^ (Supplementary Figure 2B).

We sought to understand how the “evolved” T6SS cores are distributed taxonomically, in order to determine if there is a bias in the distribution of certain “evolved” cores by a given taxonomic group. Strikingly, we can see that “evolved” Hcp seem to be overwhelmingly restricted to *Escherichia* (Supplementary Figure 3A, 10 o’clock to 1 o’clock), which represents 91.4% (1702/1861) of all genomes with “evolved” Hcp. This refines our previously mentioned observation: “evolved” Hcp is scarce, perhaps because it is narrowly distributed taxonomically, and likely appeared only once in an *Escherichia* ancestor during microbial evolution. *Salmonella* species seem to have a strong bias towards using mostly “evolved” PAAR (Supplementary Figure 3A, 1 o’clock to 2 o’clock). Interestingly, *Vibrio* is split into two subpopulations: one that largely uses “evolved” VgrG, and another subpopulation that largely uses “evolved” PAAR (Supplementary Figure 3A, 4 o’clock to 5 o’clock). The subpopulation that uses mostly “evolved” VgrG is largely made up of *V. cholerae* (83% of those with “evolved” VgrG only), and the subpopulation that encodes mostly for “evolved” PAAR is largely made up of *V. parahaemolyticus* (60.78%). We speculate these taxonomic biases may have to do with the fact that these effectors have an N-terminal core domain that needs to be compatible with the underlying T6SS core machinery in order to be loaded and fired.

We then asked if it was possible for a genome to encode a T6SS with strictly “evolved” cores, i.e. is it possible to build a T6SS without regular, “unevolved” cores. We found that virtually no genomes have strictly “evolved” cores (0.4%; 55/11752). Yet, 77% (9087/11752) of genomes contain at least one “evolved” core, showing their extensive utility as part of T6SS function. Looking closer, it is rare to find genomes with strictly “evolved” Hcp (0.5%; 63/11289) or strictly “evolved” VgrG (7%; 752/10663), yet it is relatively common to find genomes with all PAAR being “evolved” (27.2%; 2777/10180). This suggests that “evolved” PAAR does not affect efficiency of T6SS assembly, while “evolved” VgrG trimers and “evolved” Hcp hexamers may have some cost to T6SS functionality.

Many times bacterial genomes contain a main T6SS operon that encodes all the structural genes and effectors, both regular and “evolved” effectors. However, often there is also a genomic “orphan” or “auxilliary” operon, which contains just a small fraction of a T6SS operon, such as an effector gene and its cognate VgrG to which it non-covalently binds ^41^. We asked whether there is a preference for “evolved” T6SS cores to be encoded in the main T6SS operon, or in smaller orphan operons by counting the number of surrounding T6SS core genes. Regular Hcp with no C-terminal domain is encoded both in operons and outside of operons. In contrast, “evolved” Hcp have a strong bias for being encoded in orphan operons (98% of “evolved” Hcp have 0 or 1 nearby T6SS core gene), with a minority present in main operons (Supplementary Figure 3B). VgrG and PAAR are found in large numbers in both orphan and main operons, suggesting a lack of bias ((Supplementary Figure 3B).

Taken altogether, we conclude that “evolved” Hcp has very different characteristics as compared to “evolved” VgrG and PAAR. “Evolved” Hcp are very rare overall and appear nearly exclusively in a single genus. When “evolved” Hcp do occur, they have comparatively short C-terminal domains, likely representing Hcp-toxin fusion events. Furthermore, Hcp are mostly encoded in orphan T6SS loci, rather than in main T6SS operons, which may support the high cost of these effectors for a functional T6SS. Namely, it is plausible that if evolved Hcp were encoded within the T6SS operon there was a higher likelihood of yielding nonfunctional T6SS complexes, perhaps due to stoichiometric imbalances of “evolved” vs. “unevolved” Hcp.

### Many C-terminal T6SS core extensions have no annotation

Since T6Es act many times as antibacterial toxins^42^, exploration of C-terminal domains of “evolved” T6SS cores could provide a valuable resource for identification of novel antibiotic proteins. To classify the “evolved” core extensions, we surveyed all pfam domains ^43–45^ encoded in C-terminal domains of Hcp, VgrG, and PAAR. Overall, “evolved” Hcp has a very narrow variety of known pfam domains, as compared to VgrG and PAAR, which is somewhat expected due to the scarcity of these domains (Supplementary Figure 4). The most common Hcp C-terminal domains encode Colicin-DNAses, as has been shown previously ^12,46^ (Supplementary Figure 4). The vast majority of VgrG C-terminal domains have no annotation, possibly representing novel toxins (Supplementary Figure 4, gray). The C-terminal domains of “evolved” PAAR most frequently encode RHS repeats, which themselves are usually encoded upstream of a toxic domain ^38,39,47^. We also found the nuclease domain AHH, as expected from a previous study ^48^. We were intrigued by the unknown C-terminal domains, which represented an opportunity to discover novel T6Es.

### Identification of “evolved” T6Es using genomic signatures

We hypothesized that C-terminal extensions of T6SS cores that are not mapped to known Pfam domains may have toxic activity (Supplementary Figure 4, grey). To explore this hypothesis, we designed a pipeline to discover and validate these putative toxins (Figure 1, Materials and Methods). We searched 11,832 T6SS-encoding genomes, which yielded 43,546 cores with extended C-terminal domains. We clustered the C-termini of “evolved” cores with no known Pfam domain annotations into 2,324 unique families. Then, families with any pfam domains of known effectors, toxins, and enzymes were removed, leaving 713 families lacking functional annotations (Figure 1, Supplementary Figure 5). To search more sensitively, we used blastp and HHpred to further identify known effectors and toxin domains, leaving 182 C-terminal domain families (Figure 1). We further filtered for C-terminal families that were widespread phylogenetically and encoded next to a small gene, which we hypothesized was a cognate immunity gene, required for prevention of self-intoxication (Figure 1).

**Figure 1.**
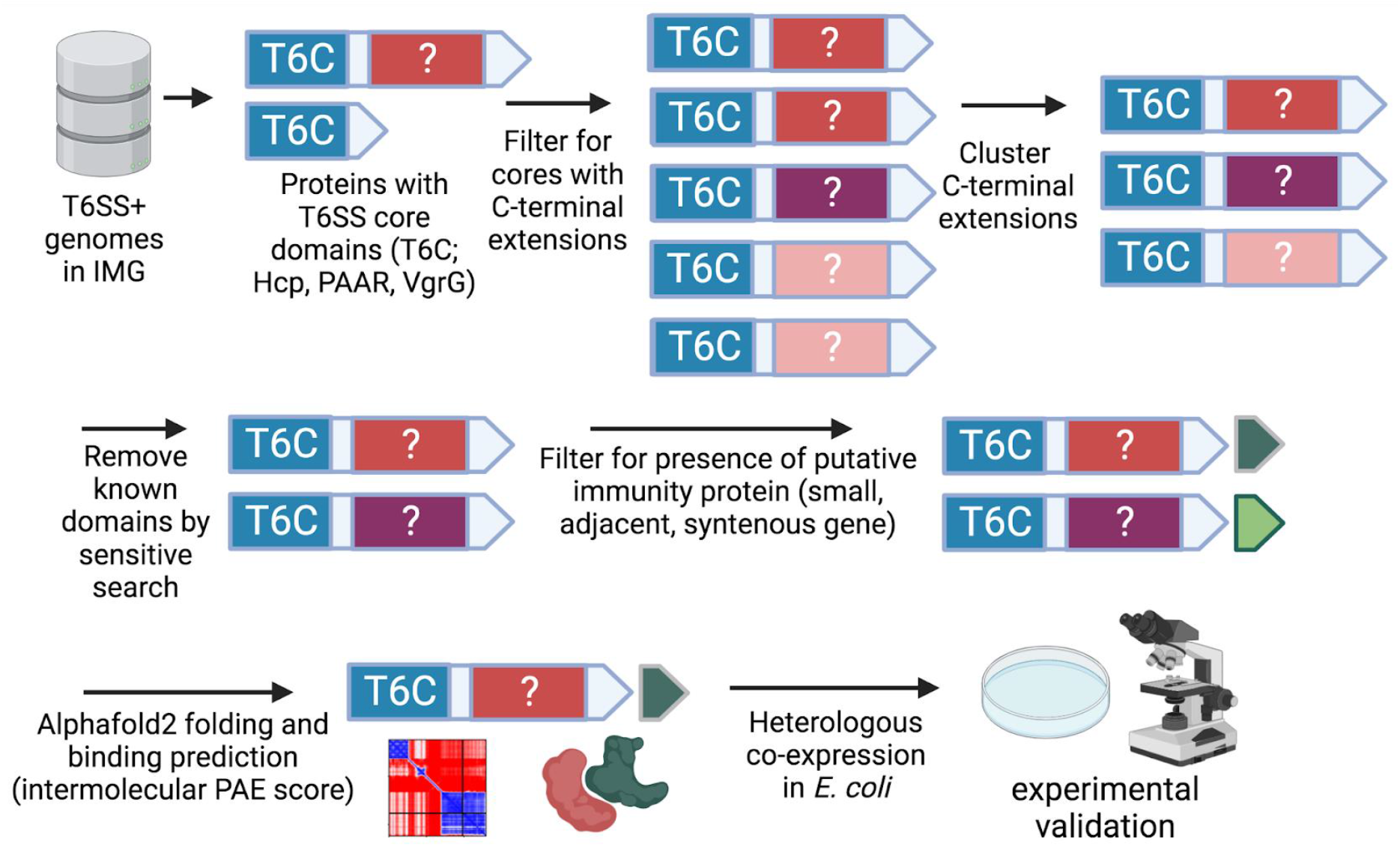
Pipeline for discovery and validation of putative T6Es and their cognate immunity proteins. T6SS-encoding genomes were searched for core pfam domains (T6Cs) (Materials and Methods), and those with C-terminal extensions were identified and considered “evolved” T6SS cores. The C-termini were clustered, and known T6E and/or toxin domains were removed. “Evolved” cores with small, syntenous genes encoded downstream were considered putative immunity genes. The pairs were tested using Alphafold2 to predict binding to one another (explained below). Those with strong genomic and structural signatures of T6Es were then tested further experimentally for toxin and antitoxin phenotypes. Created using Biorender.com

### Use of Alphafold2 for identification of predicted toxin-immunity binding

Once we had candidate T6E-immunity pairs derived from our genomic data mining, we aimed to predict whether these T6E-immunity candidates are predicted to bind one another. Since canonical T6SS immunity proteins bind to effectors and thereby neutralize their toxic activities^1^, we hypothesized that candidate effector domains and candidate immunity proteins should strongly bind one another. Weak protein-protein interactions between effector and immunity should have a high fitness cost as it can lead to unintentional self-intoxication. We took advantage of the fact that Alphafold2 can not only predict accurate three-dimensional structures of monomers, but also of multimers ^31,32,34,35,49–52^. To establish a model of protein-protein interactions using Alphafold2, we used an external dataset of other toxin-immunity pairs derived from selected T6E pairs from SecReT6 database, toxin-antitoxin pairs from TADB database, and polymorphic toxin-immunity pairs identified by our laboratory ^53–56^. We folded dimers of cognate toxin-immunity pairs, and as a control, we also folded randomized toxin-immunity pairs, i.e. a toxin from genome X and an unrelated immunity from genome Y, to evaluate the predicted interaction in a random case.

Using these data, we analyzed their predicted aligned error (PAE) scores, which is output by Alphafold2. PAE scores are used to describe the folding of one protein, but also of multimers. PAE represents the error of the positioning in space of a pair of amino acid residues (Supplementary Figure 6). In other words, a low PAE score indicates confidence in the positioning of two residues located in space relative to one another. In the case of a single protein, low PAE scores across a range of amino acid pairs results in a high confidence structure for that range of amino acid pairs (Supplementary Figure 6A-D). In the case of two proteins, low PAE scores in relevant ranges of amino acid coordinate pairs means that the two proteins are predicted to have a fixed positioning in space, i.e. they have a high confidence, stable dimer conformation, implying binding ^57^ (Supplementary Figure 6E-J). We therefore expected an overall low predicted aligned error (PAE) score distribution for a toxin-immunity pair, and an overall high PAE score distribution for a randomized pair.

In order to predict whether characteristic, high-affinity binding occurs, we input the aforementioned outside dataset of known toxin-immunity pairs (and randomized controls) into Alphafold2’s multimer mode ^31,58,59^ (using colabfold local, Materials and Methods). We observed that known effector-immunity pairs’ PAE distributions tended to have higher mean scores than those of randomized pairs’ PAE distributions (Figure 2A), suggesting we could differentiate binding vs. non-binding signals based on Alphafold2’s PAE score. To formalize and quantify the detection of binding, we trained a logistic regression classification model on these data, to classify “interacting” vs. “non-interacting” (Figure 2B-C). The model was able to separate pairs of proteins which bind one another vs. those that do not with an accuracy of 0.88, and an F1 score of 0.828 (Figure 2D).

**Figure 2.**
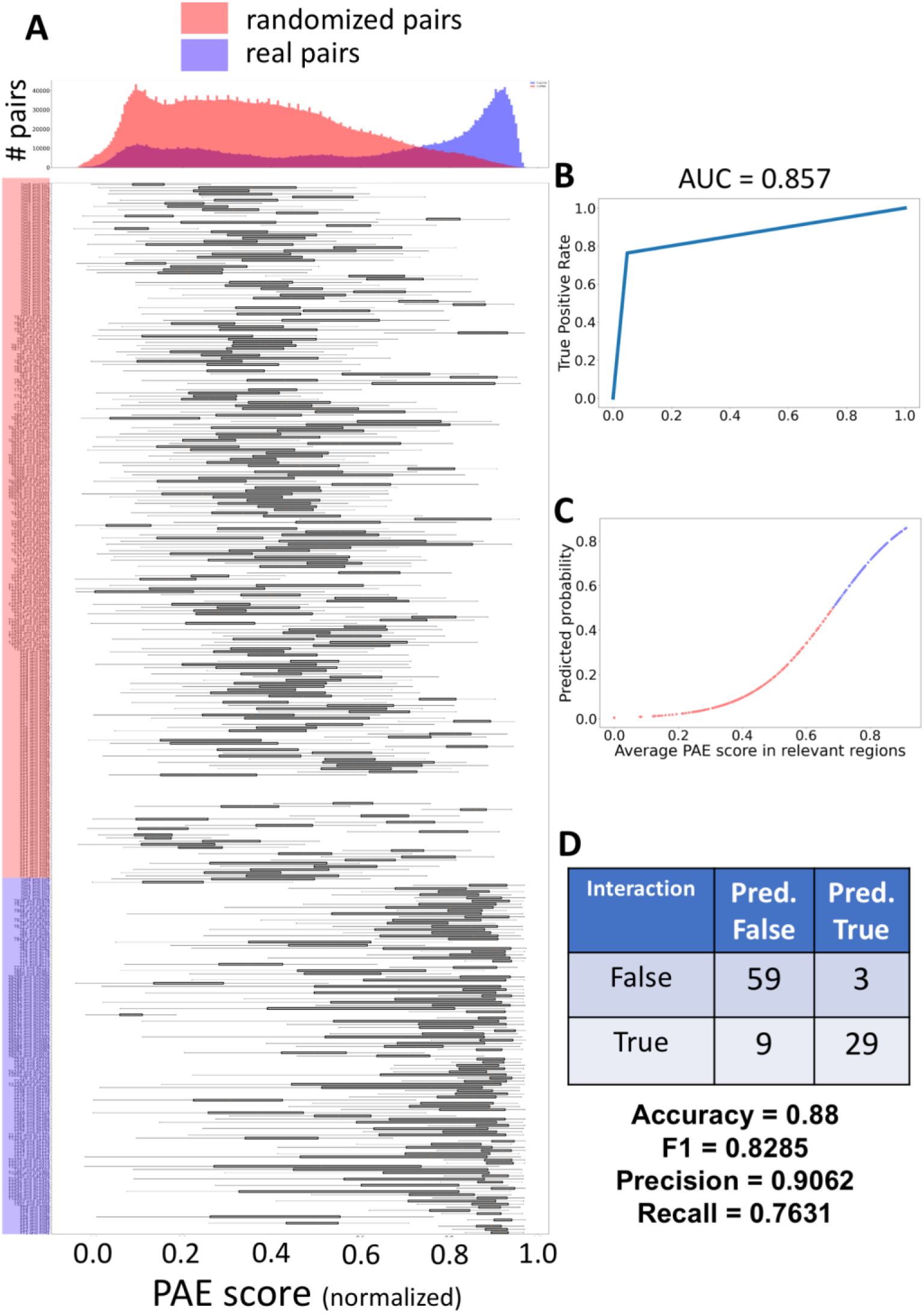
Logistic regression model of T6SS, PT, and TA toxins’ Alphafold2 PAE scores. (A) Pairs of toxin-immunity proteins were labeled as known partners with expected interactions (blue) or randomized partners with unexpected interactions (red). Each pair was run through Alphafold2-multimer, and the PAE scores between coordinates representing interprotein interaction (Materials and methods) were and plotted overall (top), or in box plots representing individual pairs (bottom). (B) ROC-AUC curve of a logistic regression model that was trained on the mean PAE scores from each pair, and classified them as “interacting” or “non-interacting”. Each pair’s probability of being “interacting” or “non-interacting” plotted, showing the logistic curve. (D) Confusion matrix of the model (top), as well as measures of model performance (bottom).

From our pipeline of 182 C-terminal domain families from “evolved” T6SS cores, we manually filtered for 15 candidates with strong genomic signatures suggesting they may indeed be T6Es (Supplementary Figure 5). We then applied the trained logistic regression model to these 15 putative T6Es and their cognate immunity genes, as well as on randomized non-cognate pairs. On this unseen test set, the model predicted 12 out of 15 pairs of putative T6E-immunity pairs as interacting (binding); 2 pairs were false negatives, and 1 pair had poor folding of the monomers, and thus could not be analyzed for binding. Furthermore, it also predicted that the randomized pairs do not bind one other, with an overall accuracy of 0.84 (Figure 3). We conclude that the genomic evidence we used to identify putative T6E-immunity pairs was strengthened by the use of Alphafold2’s prediction of binding, adding confidence that we have identified *bona fide* T6E-immunity pairs. This feature can be adopted more generally for other protein-protein interactions as well.

**Figure 3.**
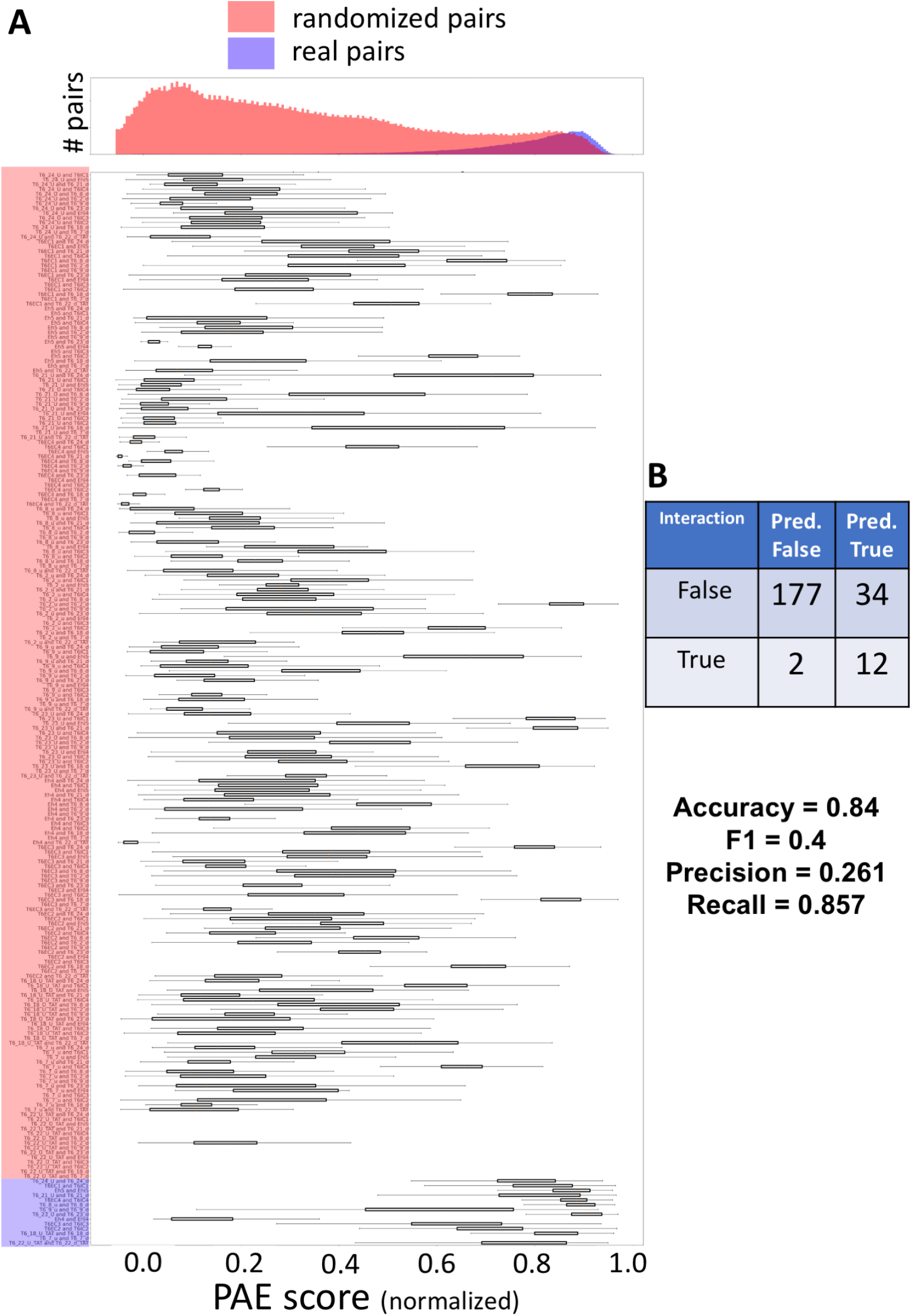
Logistic regression model applied to the test set of 15 candidate T6E-immunity pairs. (A) Pairs of T6E-immunity genes from our pipeline with expected interactions (blue) or randomized partners with unexpected interactions (red). Each pair was run through Alphafold2-multimer, and the PAE scores between coordinates representing interprotein interaction (Materials and methods) were and plotted overall (top), or in box plots representing individual pairs (bottom). (B) Confusion matrix (top), as well as measures of model performance on the test set (bottom). Note that 15 genes were input to the model but one pair folded too poorly to be evaluated by the model, so only 14 total “True” labels appear in the confusion matrix.

### Experimental validation of putative T6SS effector-immunity pairs

We hypothesized that the 15 gene pairs we predicted using genomic and/or structural signatures are indeed T6E-immunity pairs. To test this, we expressed the genes in *Escherichia coli* BL21. We expected that expression of the predicted T6Es in *E. coli* would lead to inhibition of normal growth while co-expression of the predicted T6Es and their cognate immunity genes would result in normal growth. Based on signals discussed further below, we predicted the compartment in which each putative T6E should be expressed (cytoplasm or periplasm). Localization in the periplasm was performed by the addition of an N-terminal Twin-Arginine Translocation (TAT) signal.

Upon expression of putative T6Es, 8/15 showed toxicity to *E. coli*. Furthermore, 1 of the 15 genes was synthesized but failed cloning, possibly due to high levels of toxicity. 6/15 were not toxic to *E. coli*, suggesting the prediction was inaccurate, the compartment of expression was not correct, or that the T6E is possibly targeting a cellular component not present in *E. coli* (either prokaryotic or eukaryotic). The eight putative T6Es that showed a toxic phenotype were then co-expressed with their cognate immunity proteins, and half (4/8) were saved from a toxic phenotype (Figure 4, Supplementary Figure 5, Supplementary Table 1). We named these four putative T6E-immunity pairs T6EC and T6IC 1-4 (for Type VI Effector Candidate and Type VI Immunity Candidate, respectively) (Figure 4).

**Figure 4.**
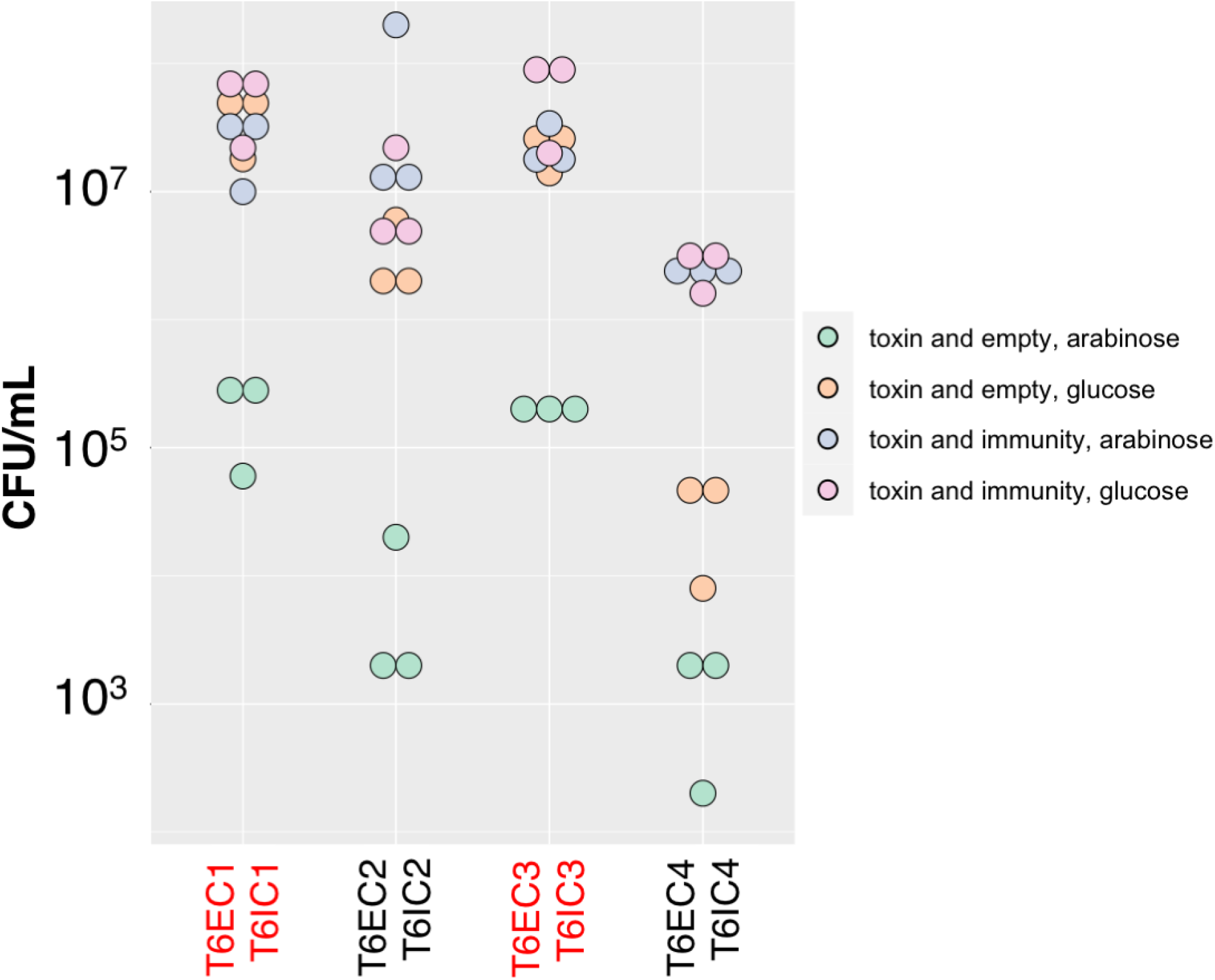
Heterologous expression of putative toxins and immunity genes in *E. coli*. Quantification of a drop assay of *E. coli* BL21 heterologously expressing T6EC1-4 in pBAD24 (arabinose induction) and T6IC1-4 in pET29b (IPTG induction), or T6EC1-4 in pBAD24 and an empty pET29b. T6EC1-4 is either uninduced (0.01 mM IPTG, 1% Glucose) or induced (0.01 mM IPTG, 0.2% Arabinose). Red text in the label specified periplasmic expression through addition of an N-terminal TAT signal sequence, except T6IC3 that has a naturally-occurring periplasmic trafficking signal.

### Structural similarity and imaging analysis suggest that the putative T6Es target bacterial DNA and peptidoglycan

Since most T6Es target various molecules required for bacterial growth^60^, we used structural bioinformatics and microscopy to determine if there are indications as to what the target in the prey cell may be for T6EC1-4.

T6EC1 is encoded in *Yersinia pseudotuberculosis* IP32881, a species that causes foodborne illness in humans^62^. T6EC1 is encoded in a T6SS operon, indicating its likely role as a T6E (Figure 5A). The protein is an “evolved” PAAR, with an N-terminal PAAR domain, and a C-terminal DUF3289 domain. A recent study of PAAR proteins also predicted this domain to serve as a toxin^48^, and we showed experimentally that it is indeed toxic to *E. coli* (Figure 4, Supplementary Figure 7). Its immunity gene, T6IC1, is encoded directly downstream of it, and contains a DUF943 domain (Figure 5A). DUF3289 and DUF943 pairs display synteny across various genomes, with variable tandem duplication of DUF943-encoding genes, in other *Yersinia*, *Enterobacter*, *Aeromonas*, and *Xenorhabdus* genomes, among others (Supplementary Figure 8). Because the T6IC1 putative immunity gene was predicted to encode a transmembrane anchoring domain, we hypothesized that the activity of T6EC1 is in the periplasm.

**Figure 5.**
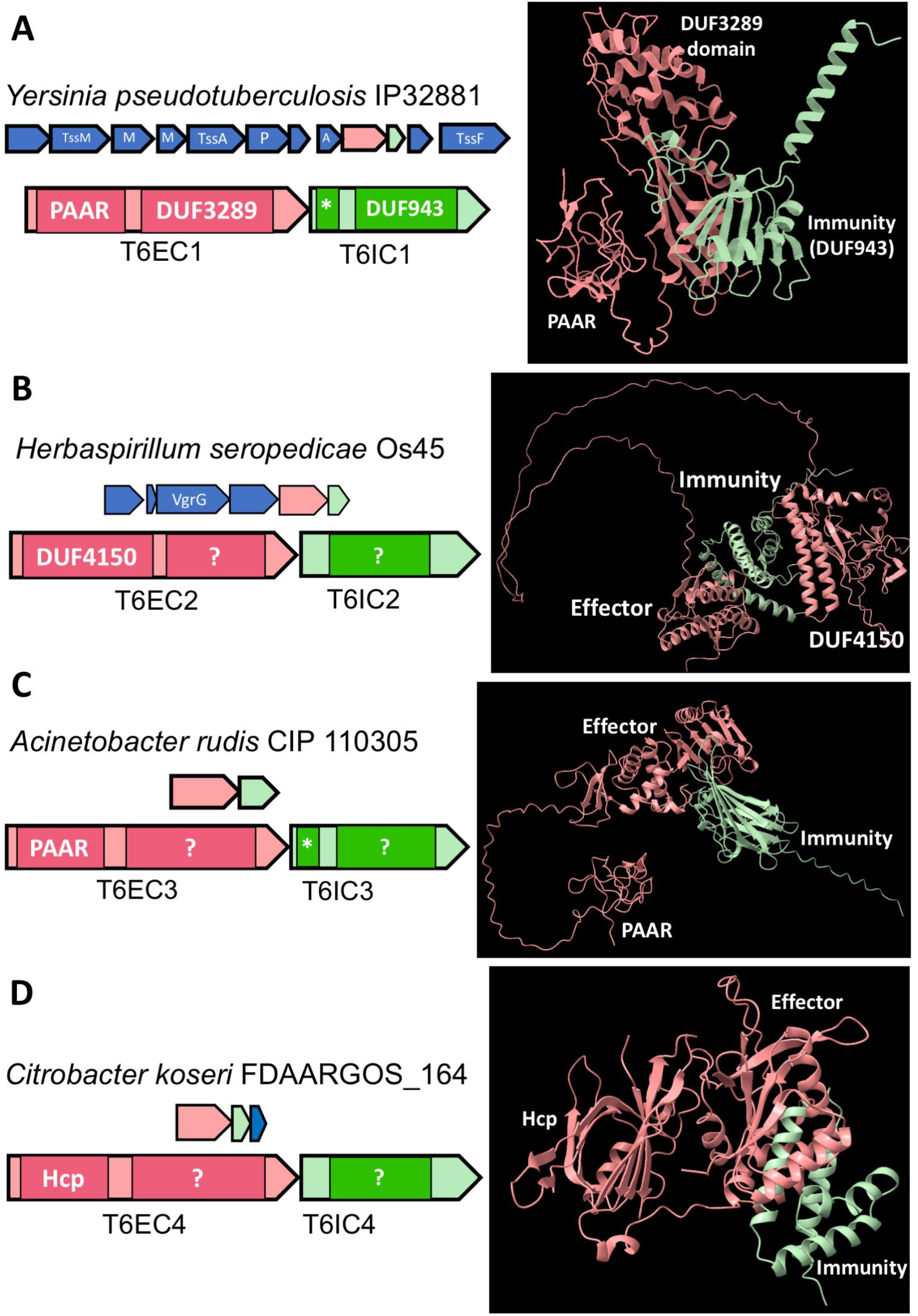
T6EC1-4 and T6IC1-4 are putative T6E-immunity pairs. (A) T6EC1 and T6IC1 are encoded in a T6SS operon (top; M = TssM, P = PAAR, A = TssA). T6IC1 has an N-terminal transmembrane domain signal, marked with an asterisk (bottom). Alphafold2 predicted structures are shown (right). (B) T6EC2 and T6IC2 are in an orphan operon containing a VgrG gene (top). (C) T6IC3 has an N-terminal signal sequence strongly suggesting it is localized to the periplasm. (D) T6EC4 and T6IC4 are in an orphan operon (top). Protein structures were visualized on ChimeraX^61^.

T6EC1 had no known functional annotation based on its sequence, so we decided to use structural searches to understand its possible activity. We searched the Alphafold2 predicted structure of T6EC1 against PDB using Foldseek, a fast, structure based search algorithm ^33,63–65^. This revealed that T6EC1 has a striking structural similarity to Colicin M (PDB: 2XTQ) (Supplementary Figure 9). This result highlights the power of structure-based searching with Foldseek, as sequenced-based annotations and search did not identify this homology, and any homologies in general.

Colicin M is known to target lipid II, the peptidoglycan precursor, by hydrolyzing it in the periplasm, thereby affecting the cell wall^66^. We reasoned that since T6EC1 was toxic in the periplasm, and has strong structural similarity to Colicin M, then T6EC1 may have a similar effect on the cell wall. To explore the possibility that T6EC1 also affects the cell wall, we expressed the toxin in *E. coli* BL21, and imaged the cells with fluorescence microscopy. We hypothesized that cells expressing T6EC1 would lose their characteristic rod shape and become round, due to loss of the protection of the cell wall against turgor pressure. By analyzing images of ∼15,000 cells we found that, indeed, a high proportion of cells expressing T6EC1 in their periplasm were round (Figure 6B-C). As a control, we showed that wild type *E. coli BL21*, as well as *E. coli BL21* co-expressing T6EC1 and T6IC1 kept their characteristic rod shape, via the analysis of ∼22,000 cells (Figure 6A, C, Supplementary Figure 10). Given these experiments, we propose that DUF3289 and DUF943 should be recognized as a putative toxin-antitoxin pair and re-annotated in interpro/pfam from its “domain of unknown function” status. Further experiments are needed to confirm the putative effector is indeed secreted in a T6SS-dependent manner.

**Figure 6.**
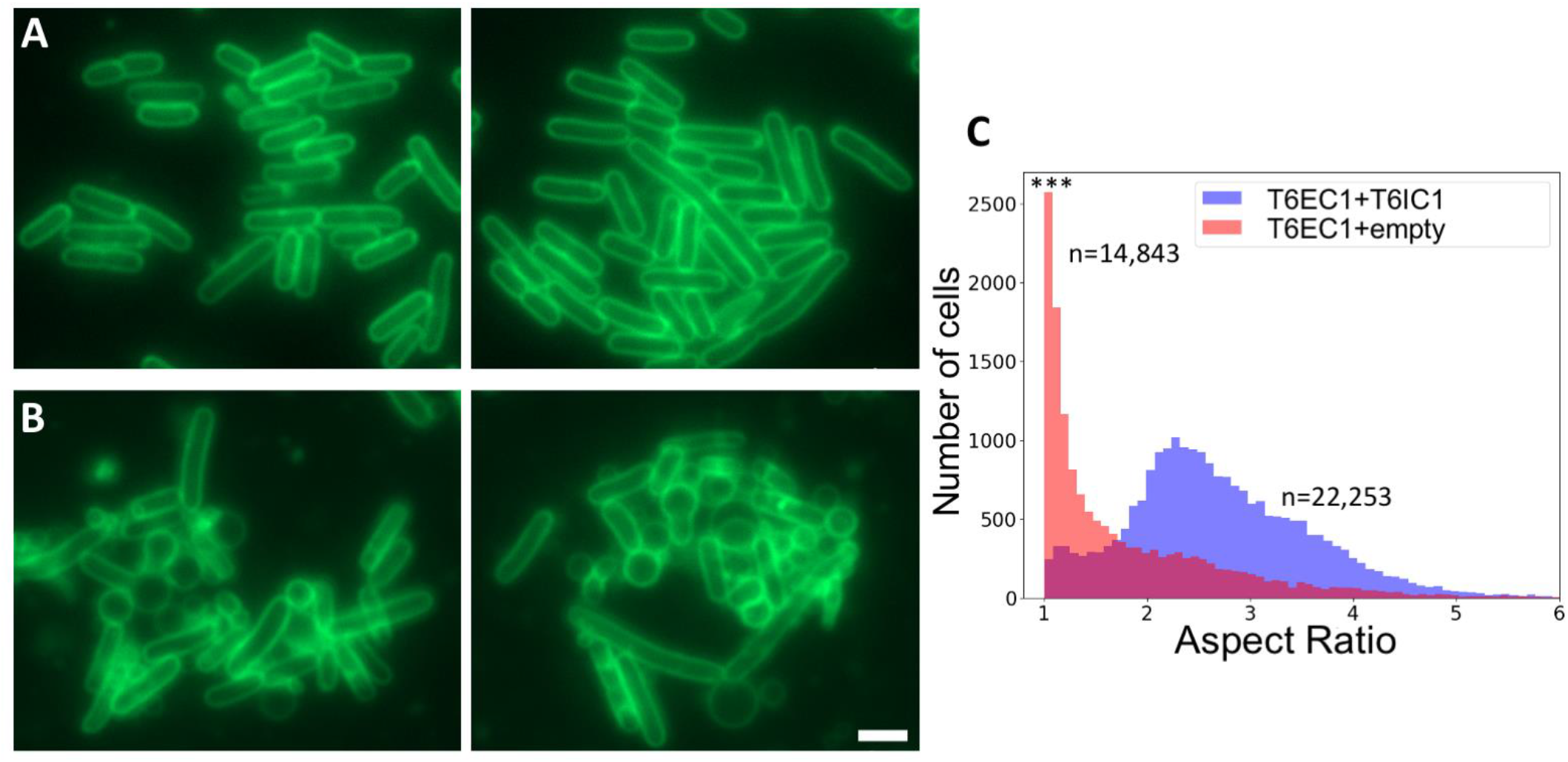
Expression of T6EC1 causes cell rounding. FM1-43 stained membranes of *E. coli BL21* expressing (A) both T6EC1 and T6IC1 or (B) T6EC1 only (empty vector control). Imaging was performed after 1h of induction of 0.01 mM IPTG and 0.2% arabinose. Scale bar = 2 micrometers. (C) Histogram of aspect ratio (a measure of length/width). Blue = both T6EC1 and T6IC1, Red = T6EC1 only, purple = overlap of histograms. An aspect ratio of 1 is a circle (equal length and width). Asterisks indicate statistical significance (Mann-Whitney U =258904636.0, pvalue=0.0).

T6EC2 is found in *Herbaspirillum seropedicae* Os45, a rice pathogen^67^. T6EC2 is encoded in an operon that, although includes a VgrG-encoding gene, does not have other T6SS genes nearby, suggesting this is a possible “orphan” T6SS operon (Figure 5B). We experimentally confirmed the toxicity to *E. coli* BL21 of the effector candidate, T6EC2, and its neutralization by the immunity gene T6IC2 (Figure 4, Supplementary Figure 11). The toxin has the structure of an “evolved” PAAR-like domain (i.e.DUF4150), with an N-terminal DUF4150 domain, and an unannotated C-terminal domain ^36,68^. Its immunity gene, T6IC2, is encoded directly downstream of it, and does not have a periplasmic localization signal, implying its activity is in the cytoplasm. The C-terminal domain of T6EC2 and T6IC2 were syntenous in other genomes of the Burkholderiales order (Supplementary Figure 12).

Sequence-based search for T6EC2 did not find any results useful in determining the function of the protein, but Foldseek search against the PDB experimentally-derived structure database resulted in a partial alignment with a EndoMS, a restriction enzyme-like dsDNAse (PDB: 5gkj) ^63,64,69^. Foldseek search of Alphafold-predicted structures also revealed that T6EC2 had a hit to the C-terminal domain of a structure of CdiA, the contact dependent growth inhibition effector (EBI Alphafold db: AF-A0A496KRC9-F1-model_v4) (Supplementary Figure 13). The C-terminal domains of CdiA vary, but many times are nucleases ^70^. Taking these two Foldseek “hits” into account, we hypothesized that T6EC2 may degrade nucleic acids, leading to cell death. We imaged *E. coli BL21* expressing T6EC2 in their cytoplasm after one hour, but we did not see any sign of damage to the DAPI signal, as we had expected. We did, however, observed extreme filamentation of cells suggestive of a severe cell division defect which may result from the biochemical activity of T6EC2 (Figure 7). In contrast, *E. coli BL21* expressing empty pBAD and pET did not have a preponderance of these long, filamented cells (Figure 7).

**Figure 7.**
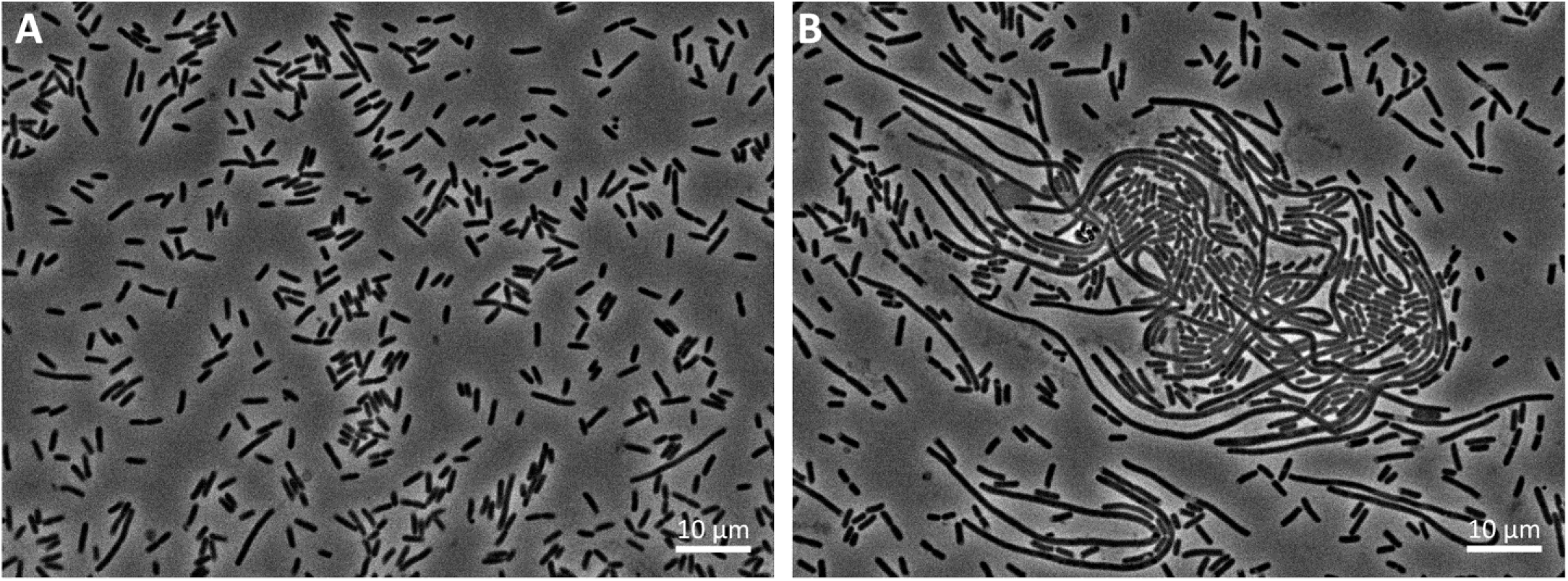
Expression of T6EC2 results in filamentation of E. coli. *E. coli* with an empty pBAD24 and pET29b vectors (A) or pBAD24 with T6EC2 and an empty pET29b vector (B) was imaged after 1 hour of incubation with IPTG and arabinose.

The toxin T6EC3 and its immunity protein T6IC3 is encoded in *Acinetobacter rudis* CIP 110305, a species that was isolated from raw milk^71^. The genes are alone, with no other T6SS genes present, which represents a possible orphan operon (Figure 5C). T6EC3 has an N-terminal PAAR domain, and a C-terminal domain with no annotation (Figure 5C). Sequence-based search did not identify the possible function of this toxin. Using signalP 6.0^72^, we found that the immunity protein T6IC3 has a predicted periplasmic localization signal, suggesting that the toxin has activity in the periplasm, perhaps against the cell wall. Furthermore, the only homolog of T6EC3 in our database was a multidomain “evolved” T6SS core encoded in a *Vibrio* species with an N-terminal VgrG domain, and also a short peptidoglycan binding domain suggesting that the toxin manipulates peptidoglycan (Supplementary Figure 15). Foldseek searches revealed a partial structural homology of the C-terminal domain of T6EC3 and SleB, a hydrolase that cleaves a specialized form of peptidoglycan during *Bacillus* spore germination (PDB: 4FET) (Supplementary Figure 16)^73^. Altogether, we concluded that T6EC3 is indeed involved in T6SS and also that it may target the cell wall. We performed microscopy to explore the hypothesis that T6EC3 will cause cell rounding. *E. coli BL21* expressing T6EC3 for thirty minutes showed many cells that had rounded (Figure 8B).

**Figure 8.**
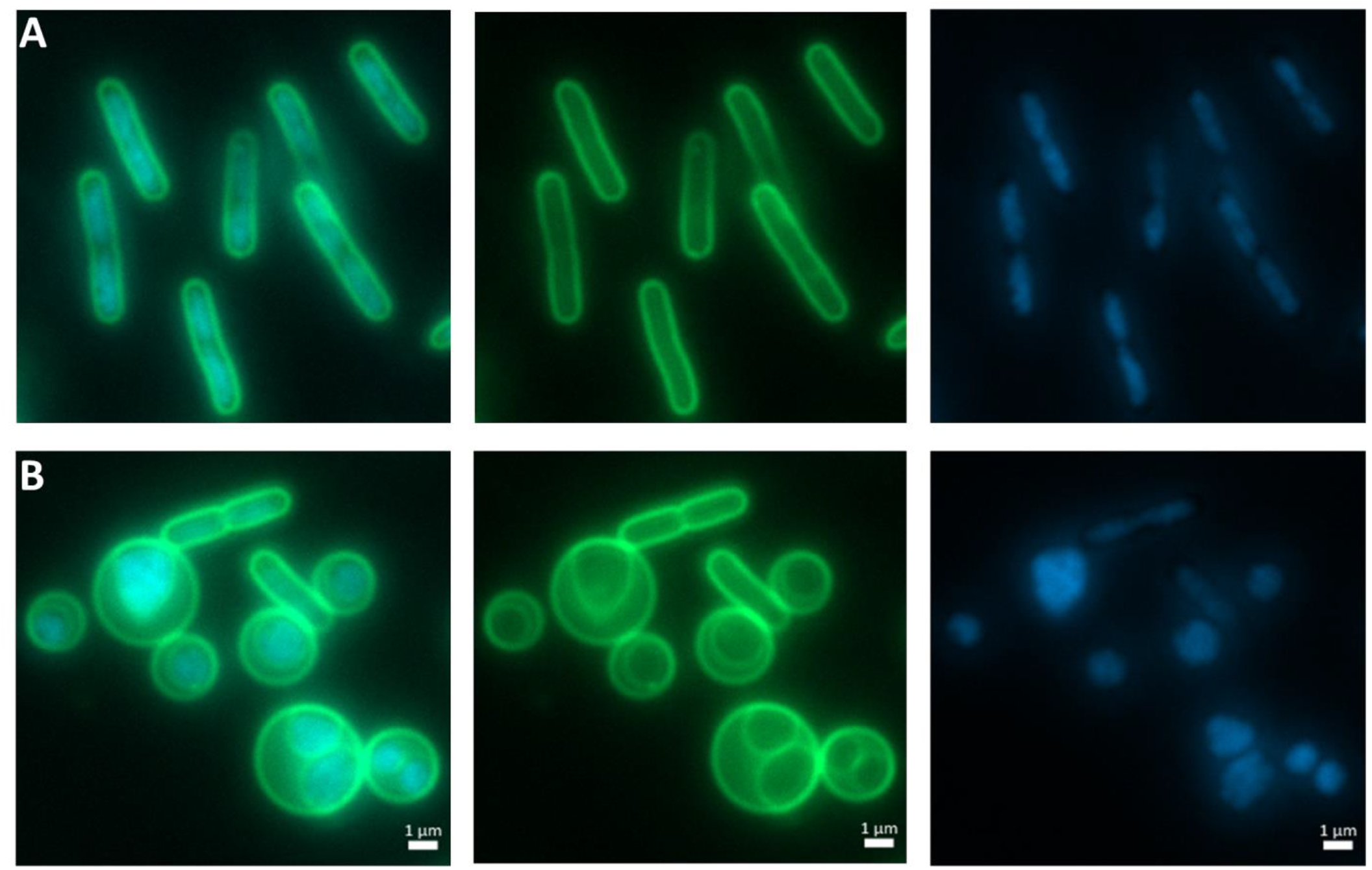
T6EC3 expression in *E. coli* cytoplasm causes rounding of cells. (A) *E. coli* BL21 expressing pBAD24 and pET29b empty vectors or (B) pBAD24 with T6EC3 and pET29b with T6IC3 after 15 minutes of induction with 0.01 mM IPTG and 0.2% Arabinose. Note the immunity protein is expressed but does not cause full saving activity (see Supplementary Figure 17).

However, in contrast to the cell rounding seen that was caused by T6EC1, the rounding caused by T6EC3 expression had peculiar double membrane structures, with DAPI signal in the inner membrane (Figure 8B). Expression of T6IC3 clearly rescued growth (Supplementary Figure 14) but could not fully inhibit T6EC3-mediated cell rounding in the microscopy conditions (Supplementary Figure 17A). Given the wealth of different mechanisms bacteria use to disrupt the cell wall^74^, we expected that other T6Es may act similarly. However, to the best of our knowledge, other studies that imaged T6Es that affect the cell wall showed cell rounding without separation of the membranes ^25,75,76^, or elongated and/or “inflated” cells that burst ^77,78^, or simply bursting ^79^. Using time-lapse imaging, we observed rod-like bacteria that lost their shape and became rounded, as well as rounded cells that eventually burst (Supplementary Movie 1). We speculate that this unique phenotype may reveal a new T6E cell wall degradation mechanism that biophysically leads to separation of the inner and outer membranes.

T6EC4 is encoded in *Citrobacter koseri* FDAARGOS_164, a medically important bacteria that was isolated from a urine catheter^80^. It is not encoded directly in a T6SS operon, but is nearby (∼20,000 bp) to a cluster of VgrG and PAAR genes (Supplementary Figure 18). T6EC4 has a rare N-terminal Hcp domain and a C-terminal unannotated domain (Figure 5D). T6IC4, its immunity gene, does not have a periplasmic localization signal, suggesting T6EC4 is active in the cytoplasm. T6EC4 and T6IC4 demonstrate synteny in various other genomes in the *Salmonella, Photorhabdus,* and *Ralstonia* genera (Supplementary Figure 19). T6EC4 is toxic to *E. coli BL21* even when expression is not induced (Figure 4, Supplementary Figure 20), showing that leaky expression is enough to stop bacterial growth. Upon induction, we see an even higher level of toxicity (Figure 4). Co-expression of T6EC4 and its putative immunity protein neutralized T6EC4’s toxic activity (Figure 4), both while the effector is induced and uninduced. This suggests leaky expression of the T6IC4 plasmid promoted cell rescue, even without induction.

Foldseek searches of its structure against the PDB database of experimentally-derived crystal structures only identified partial matches to very short segments of methyltransferases, which was largely uninformative. However, Foldseek search versus the Alphafold predicted structure database reveals a high structural similarity to Tox-REase-5 domains (Pfam15648), including to a known T6E from *Pseudomonas aeruginosa* PAO1, TseT^81^ (Supplementary Figure 21) (EBI Alphafold db: AF-Q9HXA6-F1-model_v4). Pairwise Blast search of the amino acid sequence of T6EC4 and TseT revealed no sequence similarity (data not shown), highlighting the benefit of structure based searches in determining toxin function. We therefore hypothesized that T6EC4 is an endonuclease.

We imaged *E. coli BL21* with T6EC4 and T6IC4 plasmids, with induction of T6EC4 only. The T6EC4 only strain was highly toxic, and so we imaged the strain with the presence of T6IC4, which we saw causes less toxicity, even when not expressed (Figure 9). We stained the cells with DAPI to investigate the effect of T6EC4 on the nucleotides. After one hour of toxin induction, we observed that the DAPI signal was compressed into puncta and irregular shapes in the cell (Figure 9). This result is in line with the hypothesis that T6EC4 is an endonuclease.

**Figure 9.**
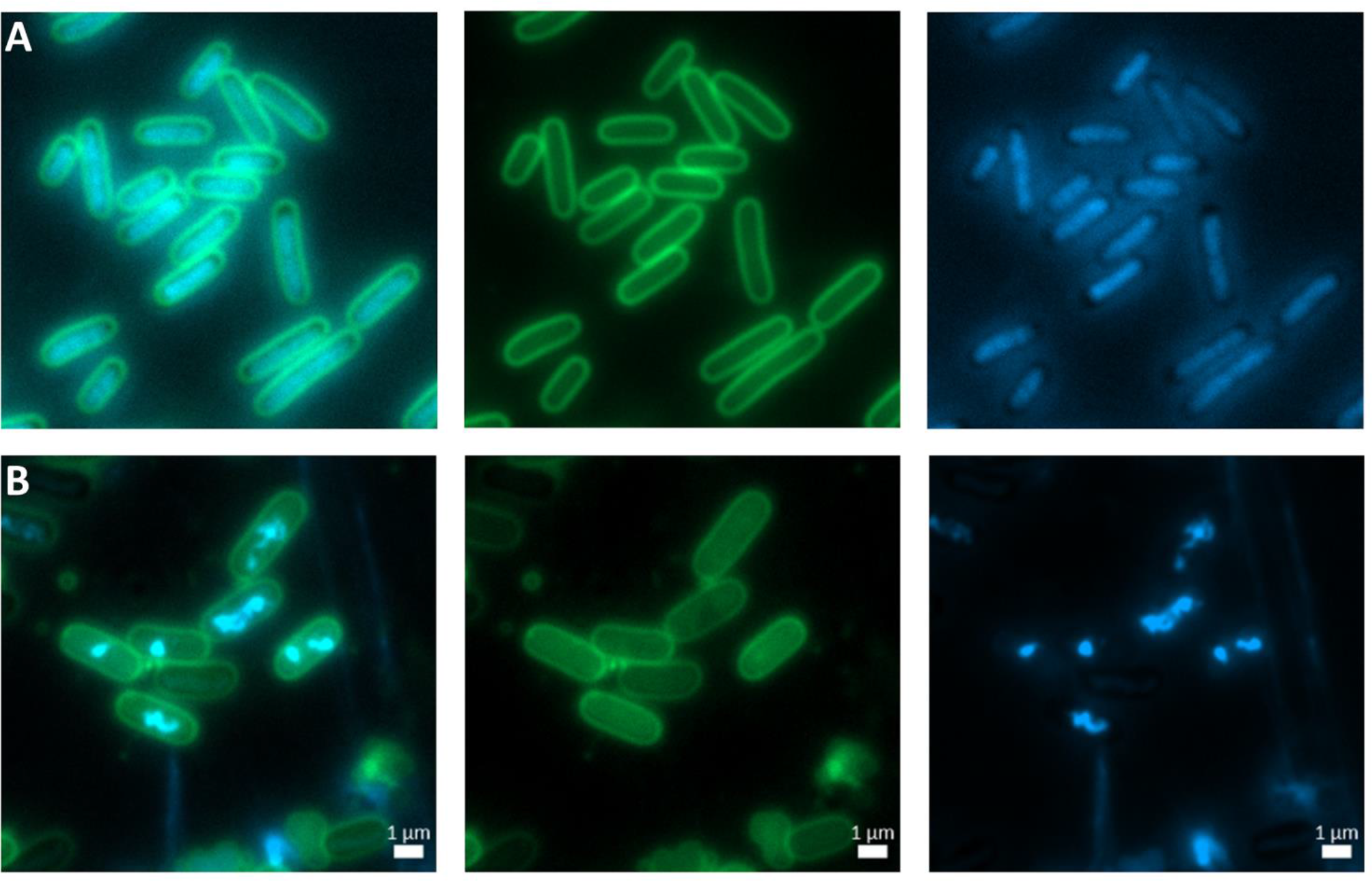
T6EC4 causes abnormalities in the DAPI signal. Empty vectors pBAD24 and pET29b or pBAD24 with T6EC4 and pET29b with T6IC4 (B) were co-expressed in *E. coli BL21* and imaged after 1 hour of 0.2% arabinose induction.

Taken altogether, we used a combined bioinformatic-experimental approach to screen and identify four putative “evolved” T6Es and their cognate immunity proteins. This is in contrast to deep, yet narrow, studies of T6Es which often result in discovery of a single T6E family from one species, albeit with more confidence. Our approach is broader, does not require a genetically tractable T6SS-encoding species, and may provide a faster first filter to screen for genes for further basic and applied research, especially given the use of novel structure-based bioinformatic tools that increase our predictive power.

## Discussion

In this study, we bioinformatically characterized “evolved” T6SS cores, and used this bioinformatic data as a starting point of a pipeline to discover novel putative T6Es. Using genomic signatures and/or Alphafold2-multimer, we identified 15 pairs of putative T6Es to study further. Using experimental validation, we showed that 8 T6Es had toxic activity (one also could not be cloned, presumably due to toxicity). Furthermore, co-expression of the candidate T6E with its putative immunity protein alleviated cells of toxicity in 4 out of 8 cases. Fluorescence microscopy was used as a first step towards investigating the possible mechanism of action of these toxins. We are aware that in order to define a gene as a *bona fide* T6E one needs to provide evidence of T6SS-dependent secretion and/or translocation, and that the presence of the gene in the attacking strain needs to correlate with higher fitness in competition assays against a prey strain that lacks the immunity protein. Despite the absence of this experiment, we highlight that an advantage of our study is that we used computational evidence combined with a simple toxin-immunity confirmation bioassay, which are both relatively straightforward and quick to perform. Another important advantage of our study design is that the organism under study does not need to be obtained/grown in the laboratory. Indeed, our putative T6Es were all identified in non-model and/or clinical isolates, making mutagenesis in these strains challenging or dangerous. Our pipeline may therefore also be applied to “evolved” cores identified in the vast number of publicly available shotgun metagenomes for discovery of novel toxins.

This study offers a parallel approach to the study of T6Es, that is meant to complement current gold standard assays. Still, we have identified multiple lines of supporting evidence that the discovered toxin-immunity pairs from this study are likely genuine T6Es: (i) The genes are taken from T6SS-encoding genomes, and are encoded in T6SS operons, orphan operons, or are near T6SS genes. (ii) The C-terminal domains are covalently attached to N-terminal T6SS core domains Hcp, VgrG, PAAR, and DUF4150, suggesting they directly bind or are loaded onto the T6SS machinery. (iii) The predicted effectors are toxic to prokaryotic cells when co-expressed with an empty plasmid, but are not toxic when co-expressed with its downstream-encoded immunity gene. (iv) The toxin and immunity proteins have predicted intermolecular interactions, therefore, they fit the classic model of neutralization of toxins by their immunity proteins. (v) The hypothesized targets of the toxins are also classic targets of T6Es. Regardless of this supporting evidence, we wrote this manuscript with caution and termed the new genes “candidate” or “putative” T6SS effectors.

The huge progress in scientists’ ability to predict protein structure from sequence biology due to AlphaFold2 and RoseTTAFold opens new doors for investigation of biology, specifically of protein-protein interactions ^49,82,83^. AlphaFold-Multimer, which we use in this paper, has been used to model and predict protein-protein interactions in eukaryotes ^84,85^ as well as in microbes ^86,87^. We have seen that certain proteins have areas of poor folding that are many times modeled as “linkers” between N-terminal T6SS core domains and C-terminal effector domains. Although the structure in the cell may be dependent on other co-binding proteins, AlphaFold2 structures do correlate with molecular dynamic simulations, suggesting that these may be true linker sequences ^57^. The use of AlphaFold2 can also be used for identifying protein targets of toxins, i.e. by folding an effector and a variety of host proteins with AlphaFold-Multimer, the target may be discovered. Because of the critical role of immunity proteins in preventing self-intoxication of cells from toxins, this is a useful training set of pairs of proteins which bind one another, and evolve rapidly, generating variety. Type II toxin-antitoxin systems, polymorphic toxin-immunity genes, and other secretion system effector-immunity pairs can also be investigated using this method. Indeed, work in this direction has already begun ^88^.

Foldseek searches provide the high-speed answer to the sudden explosion of hundreds of millions of protein sequences available due to the predicted structure databases from Deepmind and Meta ^89,90^. In our study, we saw that sequence-based searches many times did not give us insight into putative effector domain function, but structure-based searches could. This may be a method to determine the possible function of all C-terminal effector domains of evolved cores.

Overall, we view our method as an efficient and convenient way to screen for and discover a “short list” of putative T6Es for future studies. These future experiments are required to confirm that the candidate effectors are secreted in a T6SS-dependent manner and serve for intercellular competition purposes.

## Materials and Methods

Scripts are available at https://github.com/alexlevylab/T6E_discovery

### Phylogenetic tree construction

Using the IMG database of publicly available bacterial genomes ^91–93^, we searched for genomes with T6SS marker domains TssJ, TssL, and TssM (Supplementary Table 2). Because of use of these marker genes, we expected to mainly find T6SS of subtype i and ii, as TssJ, L, and M are present in those subtypes ^94^. Phylogenetic trees were constructed using a subset of universal marker genes ^95^, i.e. 29 COGs ^96^ corresponding to ribosomal proteins out of 102 COGs in Puigbò et al. ^95^. To preserve the quality of the tree, genomes missing more than one COG were dropped from analysis. COGs from each organism were aligned separately using clustal omega version 1.2.4 ^97^ with default settings. Alignments for each COG were then trimmed using TrimAl version 1.4 ^98^ using the “automated1” flag. Genomes with missing COG sequences were added as gaps into the alignments. Each trimmed alignment was then concatenated by genome, to create an amalgamated 30 COG sequence. This was inputted into VeryFastTree, version 3.0.1 ^99^ using standard settings. Trees were visualized on iTOL^100^.

### Defining “evolved” cores

An putative “evolved” effector was defined as an Hcp, VgrG or PAAR that has an extension of at least 70 amino acids downstream to the core domain (i.e. the pfam domain that defines the Hcp, VgrG, PAAR, or DUF4150). In the case of “evolved” Hcp and PAAR, there are either one or two pfam domains that define the core domain, and all were used to search for “evolved” cores (Hcp: pfam05638, PAAR (and DUF4150): pfam05488 and pfam13665; Supplementary Table 2). However, VgrG has multiple domains that define it (pfam05954, pfam06715, pfam10106, pfam13296, pfam04717; Supplementary Table 2), so the “core domain” boundaries are less clear. Many times, the conserved domains for VgrG in the COG and TIGRFAM ^101^ domains were found in a given gene, however the pfam domain was not found in that same gene (TIGR domains: TIGR01646, TIGR03361; COG domains: COG3501, COG4253, COG4540, COG3500, COG4379). Because of these issues, we therefore defined the “evolved” VgrG based on length. Using a Gaussian Mixture Model, we searched for two distributions in all VgrG based on amino acid length, “unevolved” and “evolved”. Since most of the “evolved” VgrGs having a known fused C-terminal domain were quite large, we identified a threshold between the two distributions. We used the Scikit-learn library ^102^ with the function GaussianMixture with parameters of “full” covariance. In order to define the threshold for the “evolved” VgrG, we took the mean length of the “unevolved” VgrG distribution (724aa) and added one standard deviation (72aa), which summed to 796aa length. In other words, all genes with any VgrG pfam domain that is greater than 796 amino acids in length were considered “evolved”.

### Counting “evolved” percentage

In order to see which percentage of each core is “evolved” without phylogenetic bias, we collapsed the phylogenetic tree into groups of leaves with <0.1 branch lengths and sampled one taxon each iteration from each sub-group. Then, the percentage of “evolved” cores was counted. We sampled for 10,000 iterations, and took the final average number of “evolved” cores for each core.

### Genomic pipeline

From our database of T6SS-encoding genomes (n = 11,832), we extracted genes containing Hcp, VgrG, and PAAR domains (n = 148,414). Those with C-terminal extensions greater than 100 amino acids and less than 500 amino acids were taken for further analysis (this is because most C-terminal extensions were in this range, and large outliers were not our focus in this study). The boundaries were defined by the start and end alignments of the pfam HMM profiles with each gene. This resulted in 43,546 HCP, VgrG, and PAAR that were considered “evolved”. We then clustered the C-terminal domains using CD-HIT version version 4.8.1 using parameters for >=65% identity and >= 80% coverage on both query and subject. We then submitted the representative sequences of the C-termini to the NCBI CDD search, which identifies conserved domains. We were able to label C-termini as having a known toxic enzymatic domain, or those without a known function. Those with known enzymatic/toxin domains were discarded. This left 713 clusters with putative T6SS adaptor function (or toxic effector function). An HHpred search of representatives of each family was performed against PDB70 and those proteins with hits for known T6Es or toxic enzymes were discarded ^63,103^. 182 remaining clusters were evaluated manually by searching for annotations against nr and the SecReT6^53^ database using blastp ^104^. Furthermore, genes were examined visually on IMG/M for the presence of a small syntenous gene encoded downstream (putative immunity protein) using the “Conserved Neighborhood” function^91–93^. Number of genera encoding a C-terminal cluster was noted as well, and we preferred cluster members presence in at least two genera. This resulted in sixteen candidate pairs that we considered candidate T6E-immunity pairs. We used AlphaFold2 to determine their protein structure and if they have a protein-protein interaction (see below).

### AlphaFold2

Pairs of cognate and noncognate (random) putative effectors and immunity protein amino acid sequences were folded using localcolabfold ^58,59^ version 1.4.0 with model-type AlphaFold2-multimer-v2^31,32,49^, num-recycle 10, stop-at-score 0.9, and rank multimer. The best model was used in further analyses (i.e. the rank_1 model).

The output JSON files containing the pLDDT and PAE scores were processed first to find continuous tracts of pLDDT scores above with a rolling average above 70 (window size = 30). This defines the start coordinates and end coordinates of the protein with trustworthy folding. Then we retrieved the intermolecular PAE scores corresponding to these start and end coordinates, which define rectangular matrices in the bottom left and top right quadrants of the PAE plot (Supplementary Figure 6). The PAE score presented in the figures of this study is derived by normalizing and reversing the raw PAE score by applying the formula: (30 - pae)/30, making the highest confidence scores 1, and the lowest confidence 0. Since sometimes multiple continuous tracts of pLDDT scores with a rolling average above 70 were present (i.e. multiple domains of high quality folding), we evaluated all possible combinations of intermolecular interactions, and took the domains with the maximum score. Those proteins with poor folding (no continuous tracts of pLDDT score above 70) returned no score.

The PAE score matrices in the intermolecular regions were flattened and concatenated into a simple array of PAE scores in that region. These distributions of PAE scores were retrieved from both cognate pairs of toxin-immunity proteins (from TADB, SecReT6, and PT-PIM^53–56)^ and from randomized pairings of these data into noncognate pairs (n = 333 total pairings). The means from the intermolecular PAE scores were used in a logistic regression model (implemented with sklearn.linear_model.LogisticRegression^102^, C = 0.25) to predict if the binding was “expected” (cognate pairs) or not expected (randomized pairs). 70% of the data was used for training, and 30% of the data for testing.

PDB files produced by AlphaFold2 modeling were uploaded onto the Foldseek search server^33^ (https://search.foldseek.com/). Proteins were visualized using ChimeraX ^61^.

### Drop assays

Putative T6Es were synthesized (codon optimized for *E. coli*) and cloned into plasmids (either pET29 for immunity genes, or pBAD24 for toxin genes) by Twist Bioscience (California, U.S.A.). Using heat shock, pET29 + immunity genes were transformed into *E. coli* BL21 (DE3). Transformed colonies were then used to create competent cells, which were in turn transformed with pBAD24 + toxin genes. Overnight cultures of the strains harboring the vectors of interest were grown in LB containing kanamycin and ampicillin. Cultures were normalized to 0.5 OD600 and subsequently serially ten-fold diluted. Dilutions were spotted on LB agar containing the proper selection and inducer; plates with either 1% Glucose and 0.01 mM IPTG, or 0.2% Arabinose and 0.01 mM IPTG.

### Microscopy and Image quantification

Freshly transformed colonies of *E. coli* BL21 (DE3) with pBAD24 + toxin gene, with or without pET29 + immunity gene were grown overnight in liquid cultures of LB supplemented with the appropriate antibiotic. Overnight cultures were diluted 1:100 and grown for 3 hours until the cells were in the log phase. Induction of expression of either toxin, toxin and immunity, or immunity only was performed where relevant. After incubation, 200uL of cells were centrifuged at 10,000*g, and the pellet was resuspended in 5ul of PBS with 1 mg/ml membrane stain FM1-43t (Thermo Fisher Scientific T35356) and 2 μg/ml DNA stain 4,6-diamidino2-phenylindole (DAPI; Sigma-Aldrich D9542-5MG). Microscope slides or agarose pads (for time-lapse; 1% agarose) were imaged on an Axioplan2 inverted fluorescence microscope on ZEN Blue software version 3.1 (Zeiss).

ZEN Blue or ZEN lite software (Zeiss) was used to process and export images for figures. Image quantification for some images was carried out using a deep learning package (https://github.com/noamblum/cellstats), which is based on human-in-the-loop training of the foundation model Cellpose ^105^ using microscopy images from our laboratory.

### Visualization

Figure 1 and Supplementary Figure 1 were created using BioRender (BioRender.com).

## Supplementary Information

### Movies

**Supplementary Movie 1. Time-lapse microscopy shows cells rounding and bursting.** Three scenes are shown, the first two are of the same visual field. The first only shows membrane staining, while the second shows membrane and nucleic acid stains. The third shows membrane staining only.

### Data

**Supplementary Data 1. Genomes with T6SS marker genes.** Attached supplementary data file with accession IDs and phylogeny of the 11,832 genomes with T6SS marker genes.

**Supplementary Data 2. T6SS cores gene data.** IMG gene IDs for core genes and their associated pfam domains, both core and C-terminal domains. Number of genes of T6SS genes surrounding each gene is listed. Gene architecture shows which domains make up the gene from N- to C-terminus.

**Supplementary Data 3. Alphafold2 intermolecular Binding Scores.** Normalized PAE scores are shown for pairs of cognate and non-cognate (randomized) putative T6E-immunity pairs. The amino acid coordinates in the bottom left and top right quadrants of the PAE plots that were used to determine the scores are listed. Those with “0” for the normalized PAE score did not have good quality folding overall, and could therefore not be evaluated for binding.

**Supplementary Data 4. Alphafold2 PAE and pLDDT score files.** Each file in this archive is JSON-formatted. The main contents are the maximum PAE score, PAE scores, pLDDT score, and pTM scores.

## Supplementary Tables

**Supplementary Table 1.** Predicted pairs of putative effector and immunity proteins.

| IMG gene ID | Species | Prediction | Effector toxic? Immunity saves? | Name |
| --- | --- | --- | --- | --- |
| 2685331055 | *Yersinia pseudotuberculosis*<br>IP32881 | evolved effector | effector is toxic and<br>immunity saves | T6EC1 |
| 2685331054 | *Yersinia pseudotuberculosis*<br>IP32881 | immunity |  | T6IC1 |
| 2552975056 | *Herbaspirillum seropedicae*<br>Os45 | evolved effector | effector is toxic and<br>immunity saves | T6EC2 |
| 2552975057 | *Herbaspirillum seropedicae*<br>Os45 | immunity |  | T6IC2 |
| 2566738812 | *Acinetobacter rudis* CIP<br>110305 | evolved effector | effector is toxic and<br>immunity saves | T6EC3 |
| 2566738813 | *Acinetobacter rudis* CIP<br>110305 | immunity |  | T6IC3 |
| 2666410909 | *Citrobacter koseri*<br>FDAARGOS_164 | evolved effector | effector is toxic and<br>immunity saves | T6EC4 |
| 2666410910 | *Citrobacter koseri*<br>FDAARGOS_164 | immunity |  | T6IC4 |
| 2519497843 | *Cronobacter sakazakii* ES15 | evolved effector | Effector is toxic, immunity<br>does not save | n/a |
| 2519497842 | *Cronobacter sakazakii* ES15 | immunity |  | n/a |
| 2547634204 | *Stenotrophomonas maltophilia*<br>S028 | evolved effector | Effector is toxic, immunity<br>does not save | n/a |
| 2547634203 | *Stenotrophomonas maltophilia*<br>S028 | immunity |  | n/a |
| 2617546565 | *Pseudomonas chlororaphis*<br>*chlororaphis* ATCC 9446 | evolved effector | Effector not toxic | n/a |
| 2617546567 | *Pseudomonas chlororaphis*<br>*chlororaphis* ATCC 9446 | immunity |  | n/a |
| 2545316981 | *Escherichia coli* UMEA 3805-1 | evolved effector | Effector is toxic, immunity<br>does not save | n/a |
| 2545316982 | *Escherichia coli* UMEA 3805-1 | immunity |  | n/a |
| 2649518022 | *Escherichia coli* 102132 | evolved effector | Failed synthesis | n/a |
| 2649518023 | *Escherichia coli* 102132 | immunity |  | n/a |
| 2611846572 | *Klebsiella pneumoniae* KPPR1 | evolved effector | Effector is toxic, immunity<br>does not save | n/a |
| 2611846571 | *Klebsiella pneumoniae* KPPR1 | immunity |  | n/a |
| 2588743471 | *Kluyvera ascorbata* ATCC<br>33433 | evolved effector | Effector not toxic | n/a |
| 2588743470 | *Kluyvera ascorbata* ATCC<br>33433 | immunity |  | n/a |
| 2682676947 | *Salmonella enterica arizonae*<br>63:g,z51:- So 20/20 | evolved effector | Effector not toxic | n/a |
| 2682676948 | *Salmonella enterica arizonae*<br>63:g,z51:- So 20/20 | immunity |  | n/a |
| 2677241455 | *Vibrio parahaemolyticus* S002 | evolved effector | Effector not toxic | n/a |
| 2677241456 | *Vibrio parahaemolyticus* S002 | immunity |  | n/a |
| 2611601354 | *Burkholderia pseudomallei*<br>BDD | evolved effector | Effector not toxic | n/a |
| 2611601353 | *Burkholderia pseudomallei*<br>BDD | immunity |  | n/a |
| 2705817010 | *Pseudomonas* sp. LE6C9 | evolved effector | Effector not toxic | n/a |
| 2705817011 | *Pseudomonas* sp. LE6C9 | immunity |  | n/a |

**Supplementary Table 2.** Conserved T6SS domains used in this study.

| Database | Accession | Domain name | Used for |
| --- | --- | --- | --- |
| pfam | pfam05488 | PAAR domain | search for “evolved” cores |
| pfam | pfam13665 | DUF4150 (PAAR like) | search for “evolved” cores |
| pfam | pfam05954 | Phage GPD protein (VgrG analog) | search for “evolved” cores |
| pfam | pfam06715 | gp5 repeats (VgrG component) | search for “evolved” cores |
| pfam | pfam04717 | base V (VgrG component) | search for “evolved” cores |
| pfam | pfam13296 | T6SS_VgrG | search for “evolved” cores |
| pfam | pfam10106 | DUF2345 | search for “evolved” cores |
| pfam | pfam05638 | Hcp | search for “evolved” cores |
| pfam | pfam17642 | TssD | search for “evolved” cores |
| COG | COG3455 | VasL (TssL) | Defining T6SS+ genomes |
| COG | COG3521 | TssJ | Defining T6SS+ genomes |
| COG | COG3523 | TssM | Defining T6SS+ genomes |

## Acknowledgements

The authors thank Noam Dotan for help with phylogenetic trees and scripting, and Yaara Oppenheimer-Shaanan for technical help with microscopy. AL is generously supported by the Israeli Science Foundation (Grants #1535/20, #3300/20), Alon Fellowship of the Israeli council of higher education, The Hebrew University - University of Illinois Urbana-Champaign seed grant, the Israeli Ministry of Agriculture (Grant 12-12-0002), and ICA in Israel. AMG is generously supported by the Kaete Klausner Scholarship and was supported by a scholarship from the Israeli Ministry of Aliyah and Integration.

---

**Supplementary Figure 1.**
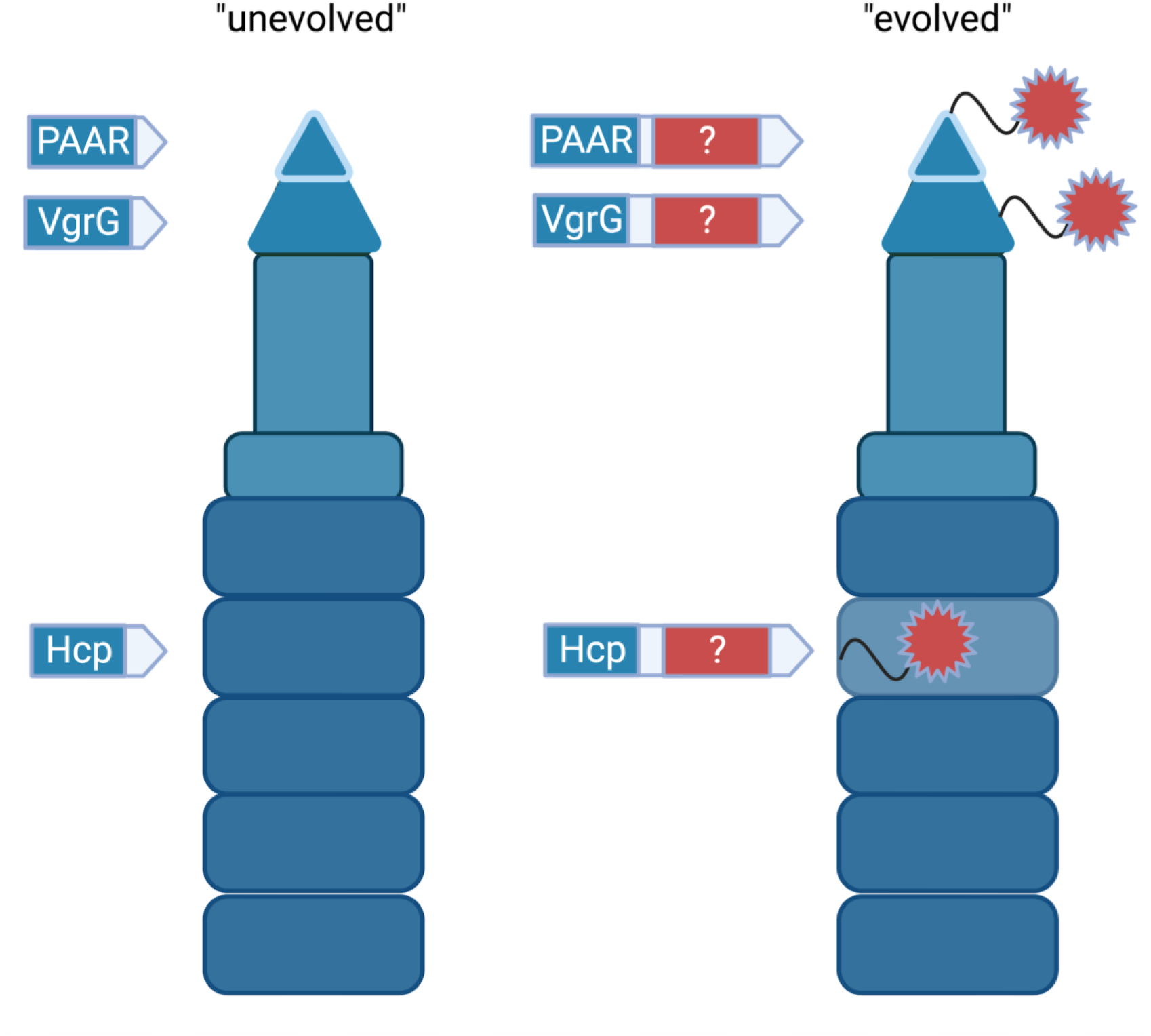
Architecture of T6SS structural proteins (“cores”) with and without C-terminal extensions (“evolved” and “unevolved”, respectively). Left: “unevolved” Hcp (tube), VgrG (spike), PAAR (spike tip). Middle: “evolved” versions of the “cores”, i.e. with C-terminal extensions. Right: a cartoon of T6SS “evolved” cores *in situ* in the T6SS machinery. Red shape represents the C-terminal domain, which many times is an enzymatic toxic domain, which is inherently loaded onto the T6SS structure, due to the N-terminal core domain. Created with Biorender.com

**Supplementary Figure 2.**
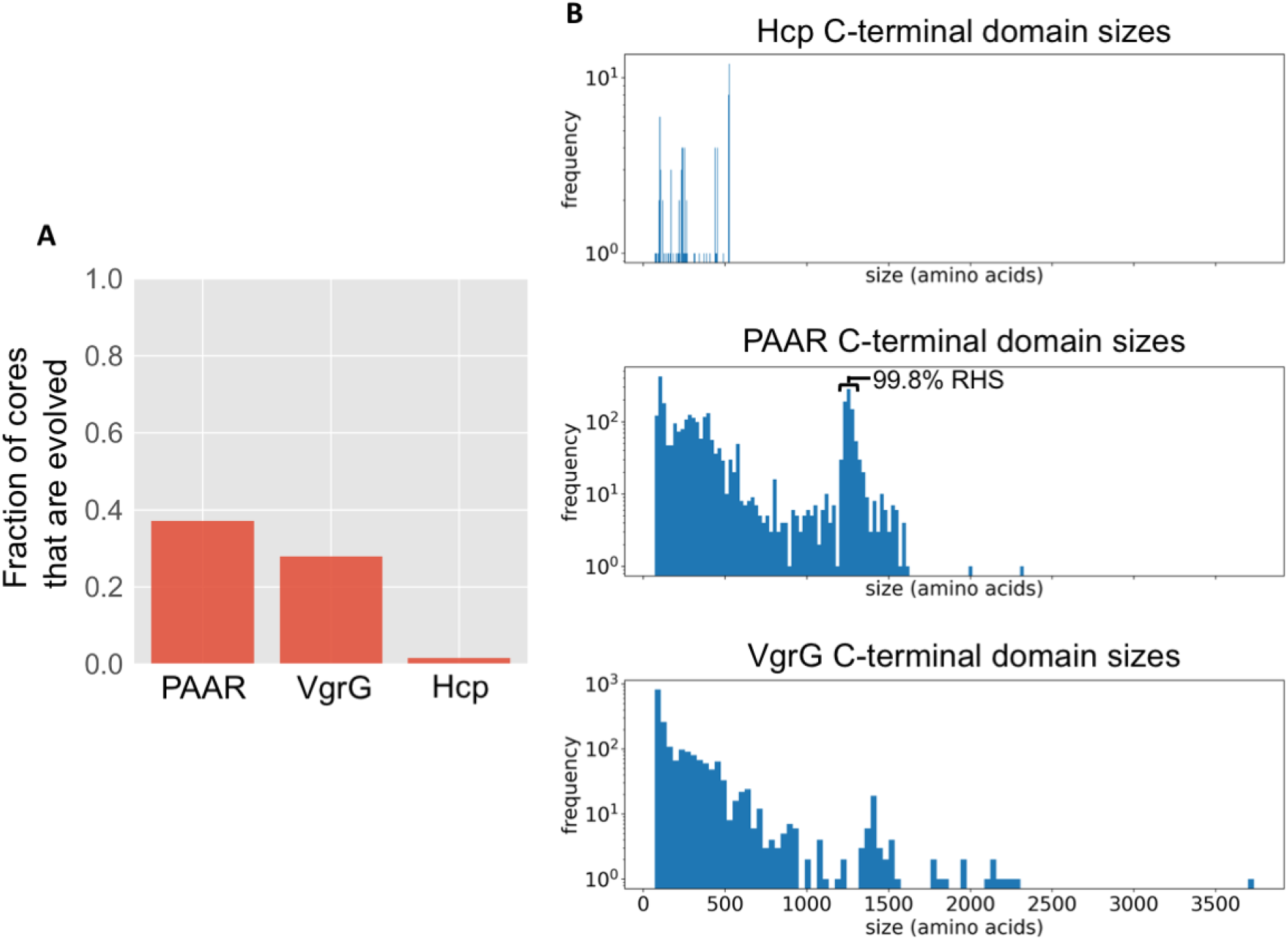
“Evolved” T6SS cores’ prevalence and C-terminal domain sizes. (A) Fraction of cores which are “evolved”. (B) Histogram of sizes of C-terminal extensions of Hcp, PAAR, and VgrG.

**Supplementary Figure 3.**
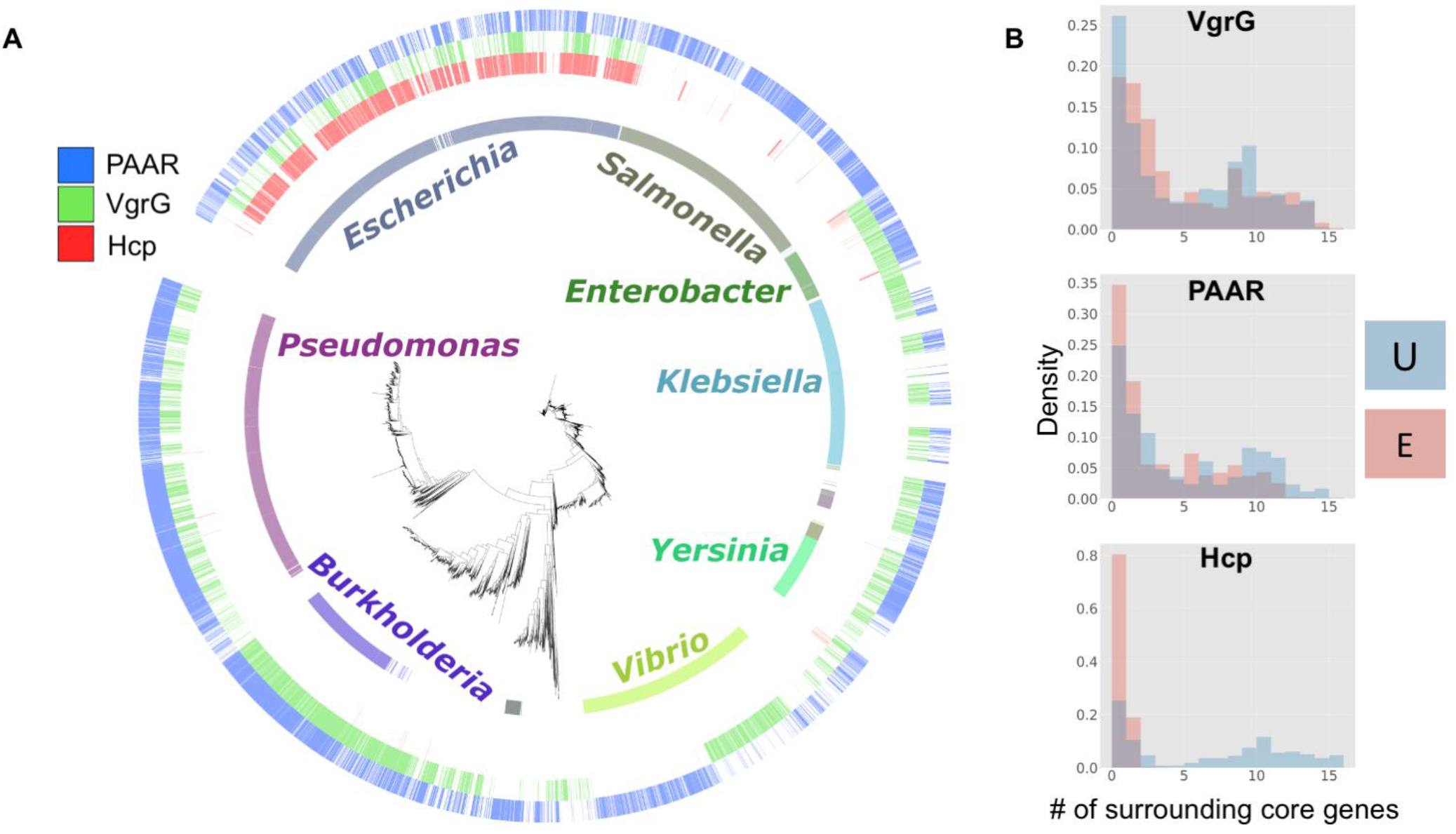
“Evolved” T6SS cores taxonomic and physical distribution. (A) Phylogenetic tree of T6SS-encoding genomes show how “evolved” PAAR, VgrG, and Hcp (outer three rings) are distributed across the Genera (inner ring). Only genera with >100 members in the dataset are shown. (B) Histograms of the number of T6SS operon genes encoded next to each core, “unevolved” (U, blue) or “evolved” (E, red). Overlapping distributions produce a purple color.

**Supplementary Figure 4.**
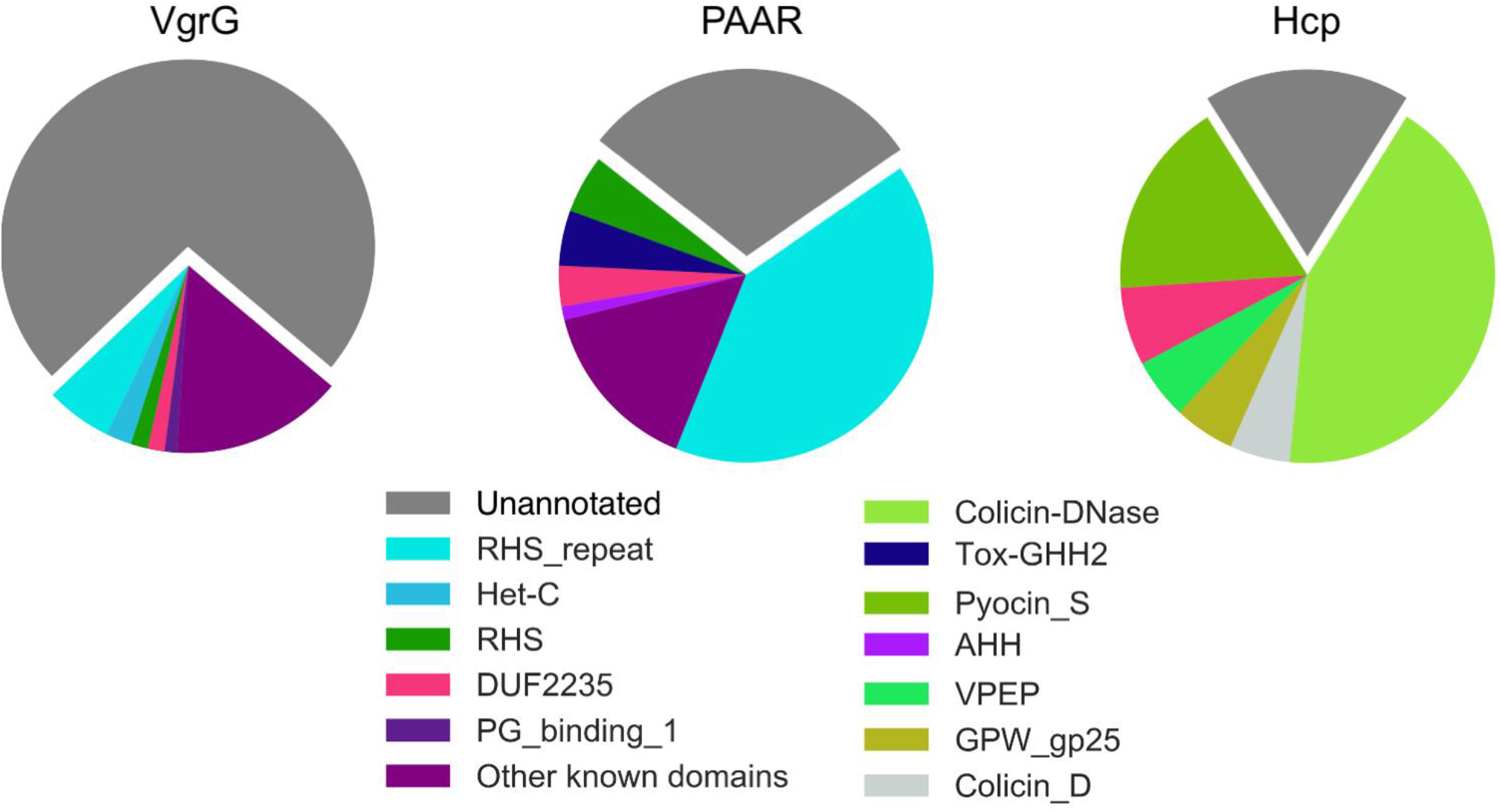
Many C-terminal T6SS core extensions have no functional annotation. Annotations of C-terminal domains were done using the Pfam database^43–45^. To avoid phylogenetic bias in plotting proportions of C-terminal domains, one genome per clade (average distances < 0.1) was sampled. This sampling was performed 100 times, each time the makeup of the C-terminal domains of “evolved” cores was noted. After the 100 iterations, the average counts of C-terminal domains were plotted here (average n ≈ 52, 576, and 21 for VgrG, PAAR, and Hcp, respectively). Domains with less than 1% occurrence on average were considered below threshold for plotting and grouped together into “Other known domains” (purple). Names in the legend are the pfam database short names.

**Supplementary Figure 5.**
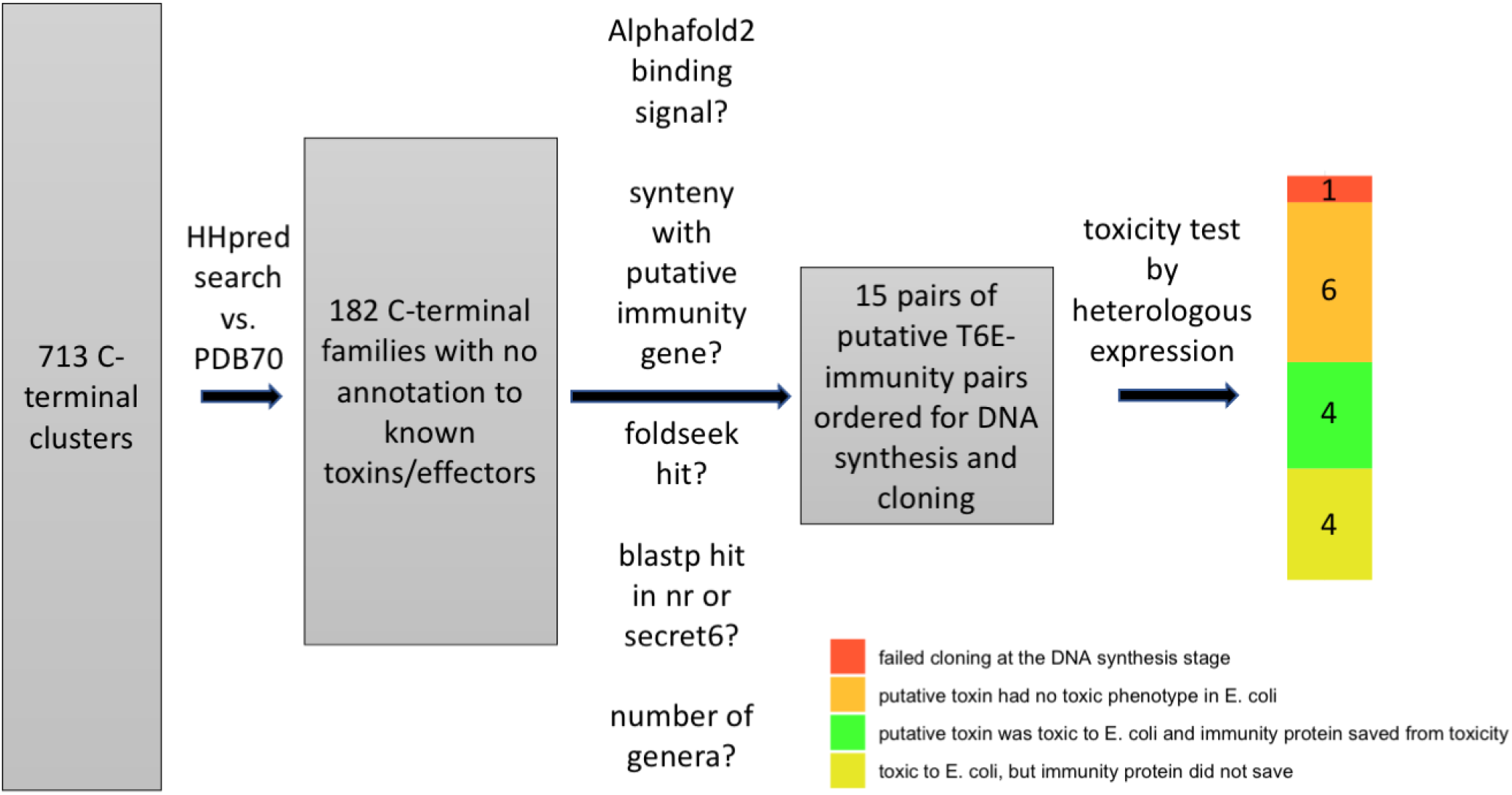
Filtered C-terminal clusters of T6Es resulted in 15 candidates for further testing. The pipeline in Figure 1 was applied to our dataset of T6SS-encoding genomes. The numbers represent the remaining C-terminal families throughout filtering steps of the pipeline.

**Supplementary Figure 6.**
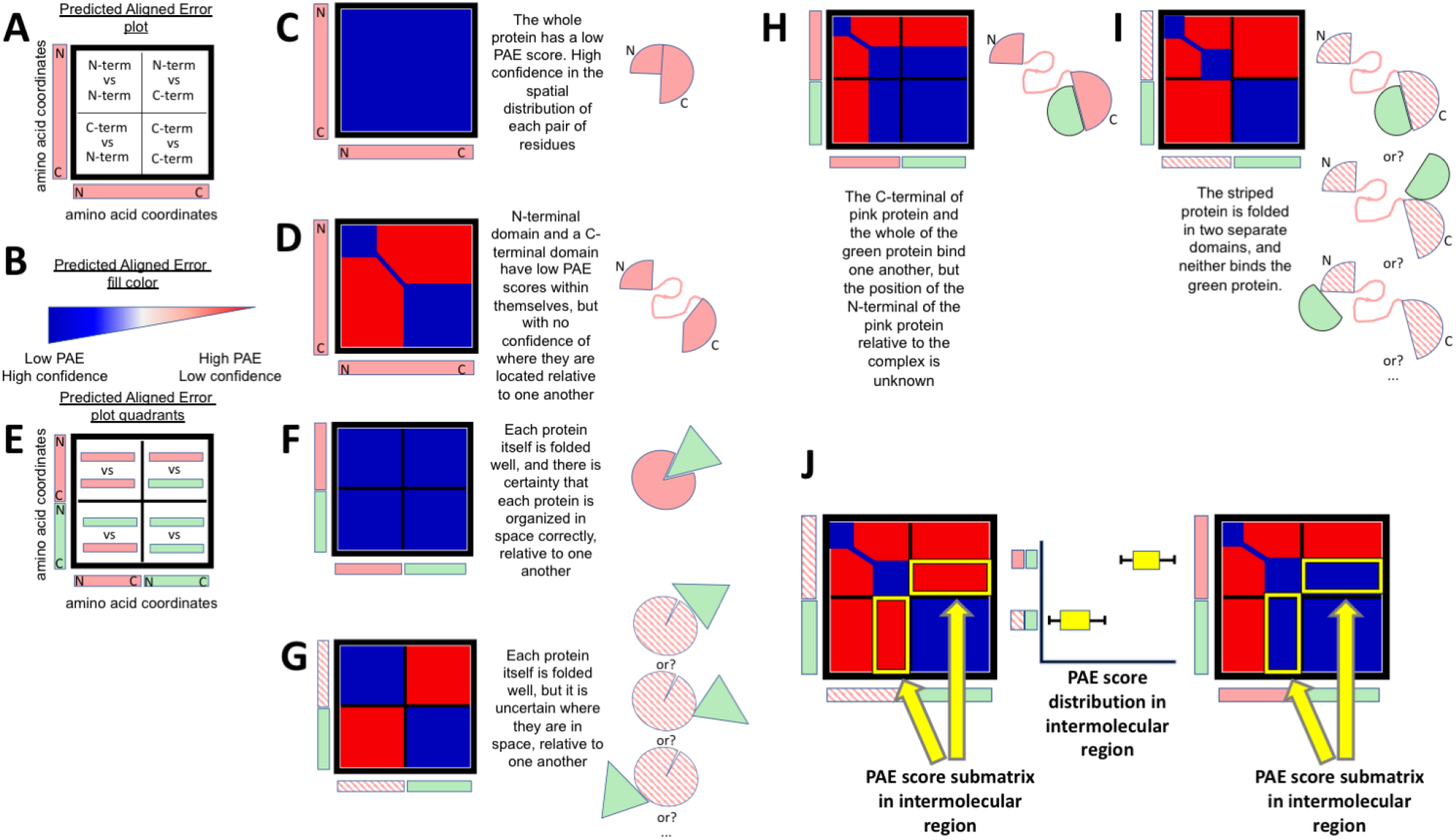
Alphafold2 predicted aligned error (PAE) plots and interpretation of single proteins. Single proteins can be folded *in silico* by Alphafold2, and among the outputs of the program are the PAE scores for each pair of residues. (A) The PAE score can be visualized as a plot where the X- and Y-axis contain the amino acid coordinates of the folded protein. Note the Y-axis’ coordinates go from top to bottom, in contrast to classic Cartesian coordinates. Each point in the graph therefore represents a pair of amino acid residues. (B) The confidence of the spatial orientation of each pair of residues is captured in the PAE score, visualized as a range of colors. (C) An ideal case of perfect folding of a single protein, where each pair of amino acids has a low PAE score, i.e. the confidence in the spatial positioning of each pair of amino acids is high. (D) A PAE plot showing two tracts where the PAE score is low, but high PAE score (low confidence) between these two tracts. This is interpreted as two domains, each with a high confidence folding, but the positioning relative to one another is unknown, and can be reasonably imagined as two globular domains attached by a flexible linker. (E) PAE score plot for multiple proteins. Two proteins of interest are folded together using Alphafold2’s multimer model. The X- and Y- coordinates are the amino acid sequences of both proteins, one concatenated onto the other. Each quadrant of the plot divides between intra- vs. inter-protein interactions. (F) An ideal PAE plot showing perfect folding of each protein alone, as well as strong interactions between the two proteins. In other words, there is high confidence in the spatial orientation of each pair of residues, including pairs on different protein chains. (G) A PAE plot showing that each protein’s folding is of low PAE, but the interaction between them is of high PAE, i.e. there is no confidence of the relative position between the two proteins. (H) A PAE plot showing a pink protein with two domains (top left quadrant), and a green protein with one well-folded domain (bottom right quadrant). The blue signal in parts of the bottom left and upper right quadrants indicate that the C-terminal of the pink protein interacts with the green protein. However, the spatial positioning of the N-terminal portion of the pink protein and the green protein has a high PAE score, i.e. low confidence. (I) PAE plot showing a striped protein with two domains (top left quadrant), and a green protein with one well-folded domain (bottom right quadrant). The red signal (high PAE) in the bottom left and upper right quadrants indicate that the orientation of the two proteins in space is unknown, i.e. they are not predicted to interact. (J) The relevant intermolecular region (submatrix) for evaluation of PAE score (yellow rectangles). The submatrix represents coordinates at which the folding of singular proteins was of high quality, but in the bottom left and top right coordinates. Since each pixel in the plot represents a single PAE score for a pair of coordinates, the submatrix can be shown as a distribution (center; box and whisker plot).

**Supplementary Figure 7.**
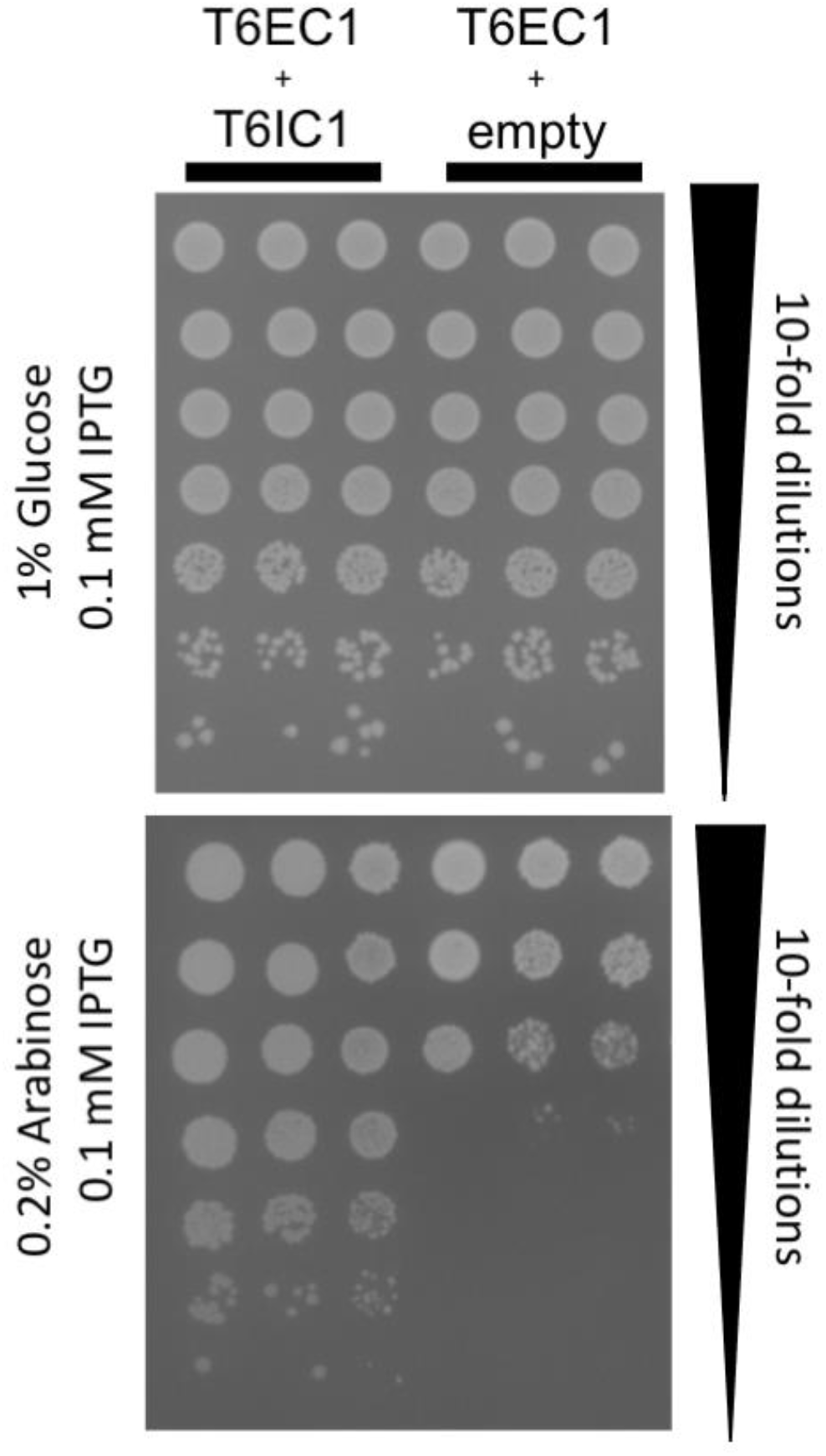
Heterologous expression of T6EC1 and T6IC1 genes in *E. coli* BL21. Drop assay of *E. coli* BL21 heterologously expressing T6EC1 in pBAD24 (arabinose induction) and T6IC1 in pET29b (IPTG induction), or T6EC1 in pBAD24 and an empty pET29b. T6EC1 is either uninduced (0.01 mM IPTG, 1% Glucose) or induced (0.01 mM IPTG, 0.2% Arabinose).

**Supplementary Figure 8.**
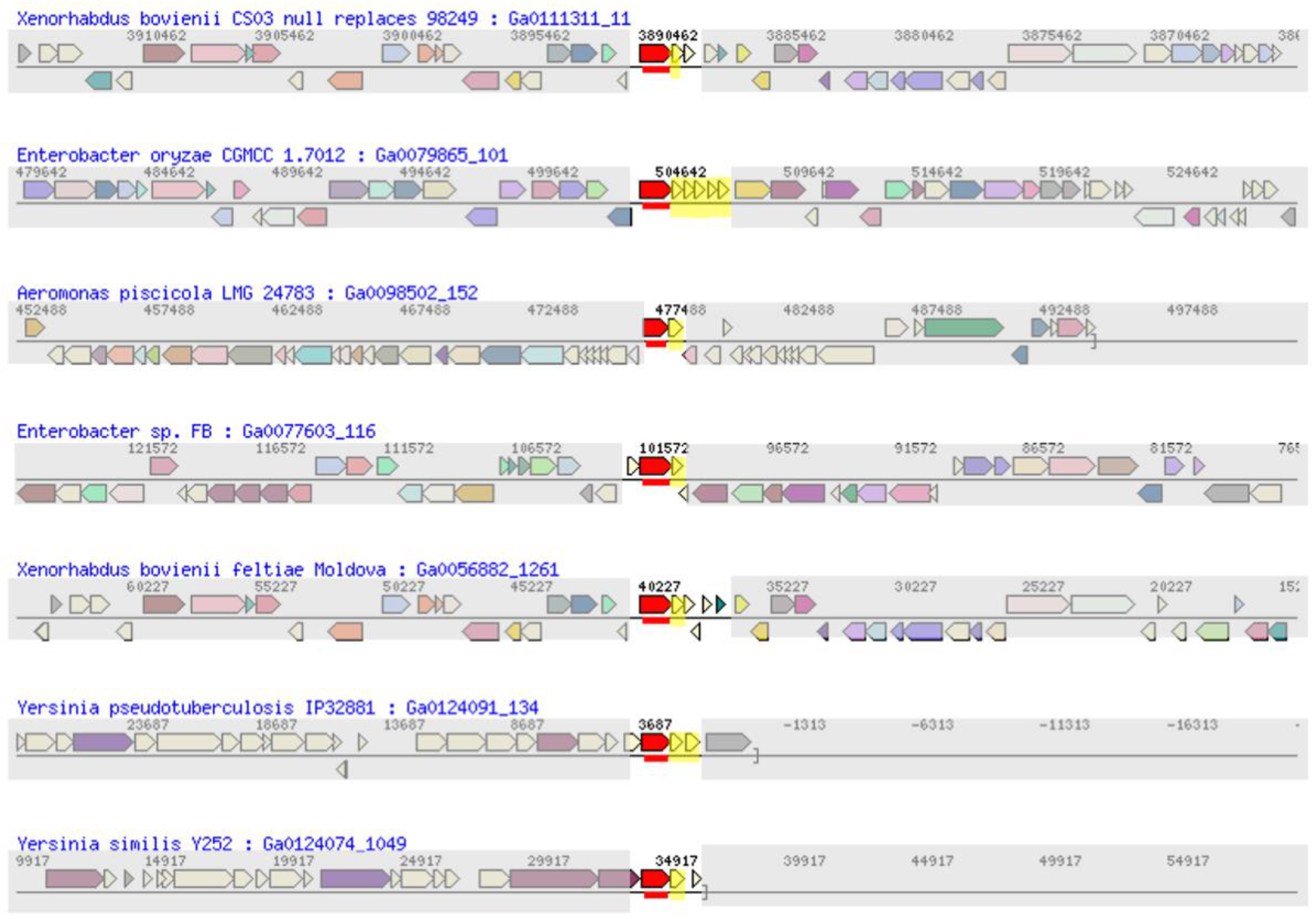
T6EC1 and T6IC1 display synteny across various genomes. Each row is a view of the noted genome (blue font). Red gene = T6EC1 and its homologs. Yellow = genes containing DUF943.

**Supplementary Figure 9.**
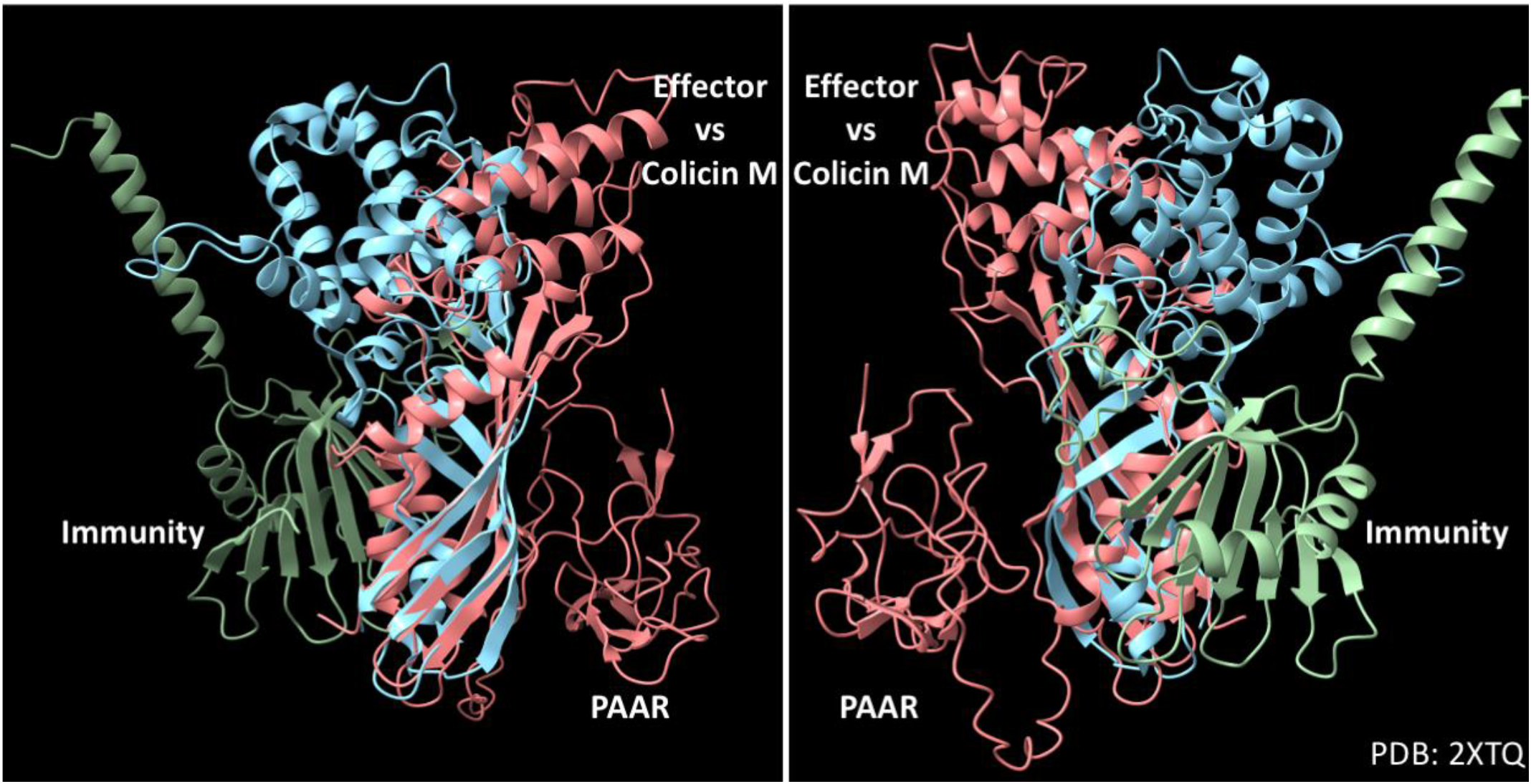
Structural search, but not sequence-based search, identified T6EC1 as a possible homolog of Colicin M. Foldseek search of T6EC1 (pink) showed a match to Colicin M (light blue; PDB accession: 2xtq). The protein structures were overlaid and visualized on ChimeraX^61^.

**Supplementary Figure 10.**
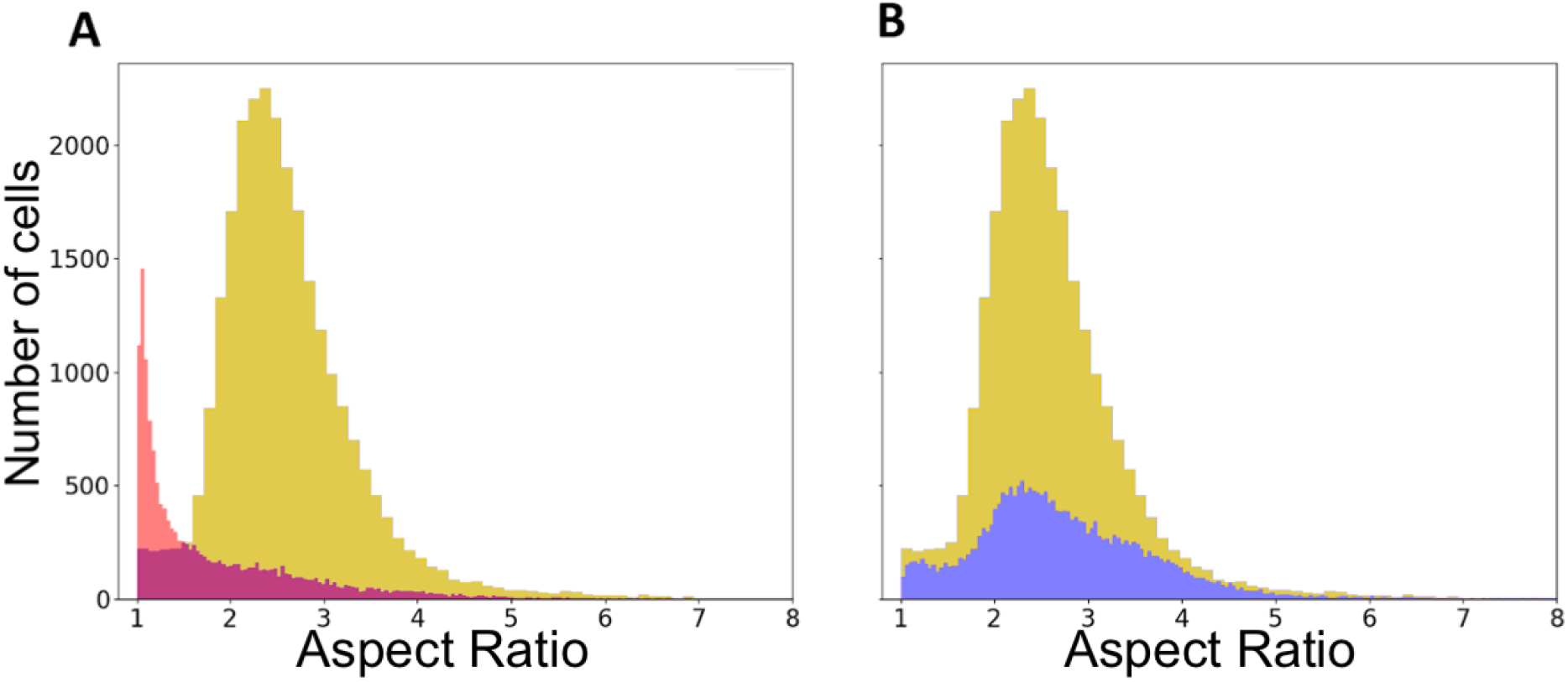
Co-expression of T6EC1 and T6IC1 results in a cell shape distribution that is similar to that of an empty vector control. (A) Histogram of aspect ratio (a measure of length/width) derived from images of FM1-43 stained membranes of *E. coli BL21* expressing (A) Red = T6EC1 only, gold = empty vectors, purple = overlap of histograms or (B) gold = empty vectors, blue = both T6EC1 and T6IC1. Imaging was performed after 1h of induction of 0.01 mM IPTG and 0.2% arabinose.

**Supplementary Figure 11.**
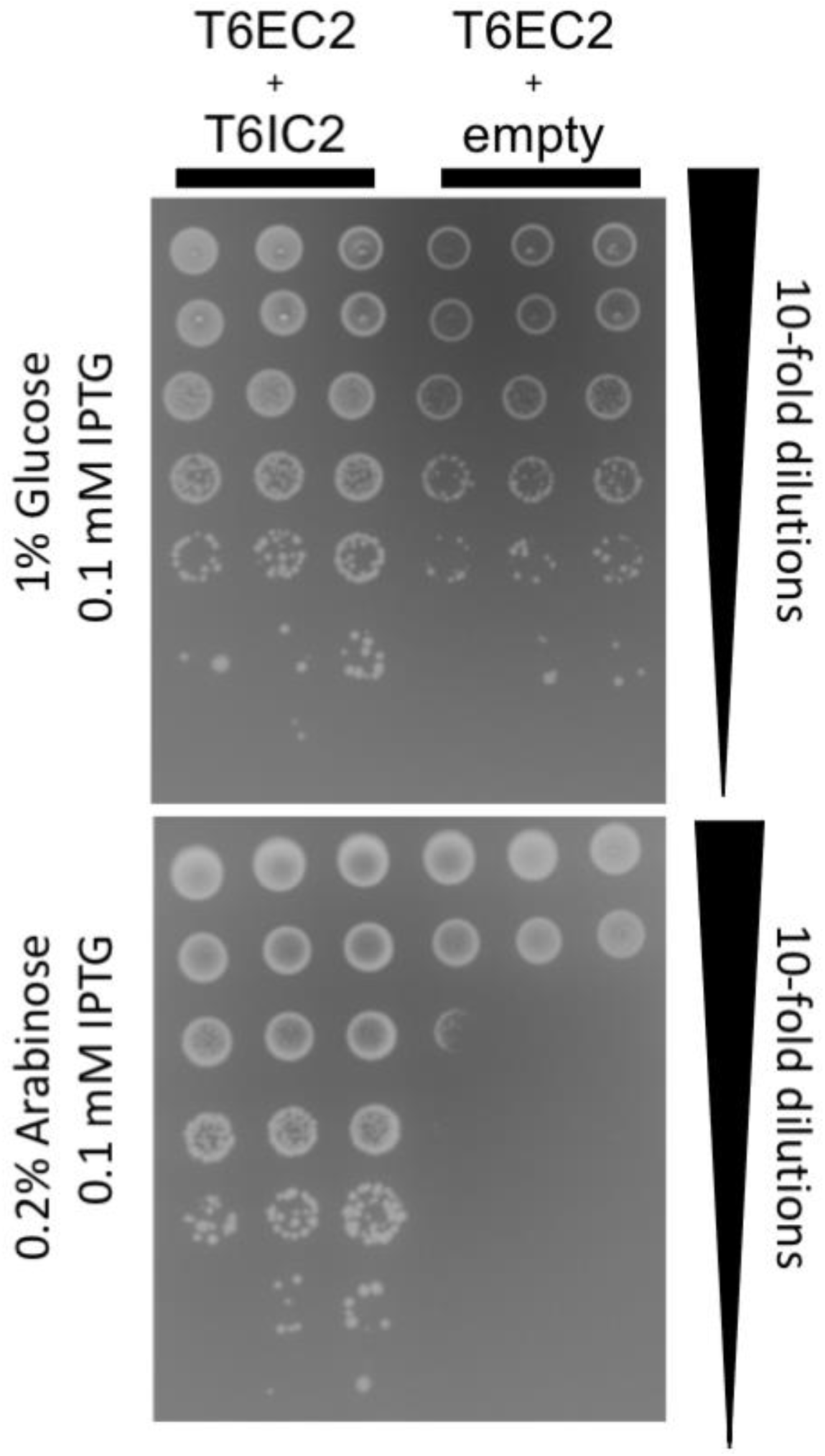
Heterologous expression of T6EC2 and T6IC2 genes in *E. coli BL21*. Drop assay of *E. coli BL21* heterologously expressing T6EC2 in pBAD24 (arabinose induction) and T6IC2 in pET29b (IPTG induction), or T6EC2 in pBAD24 and an empty pET29b. T6EC2 is either uninduced (0.01 mM IPTG, 1% Glucose) or induced (0.01 mM IPTG, 0.2% Arabinose).

**Supplementary Figure 12.**
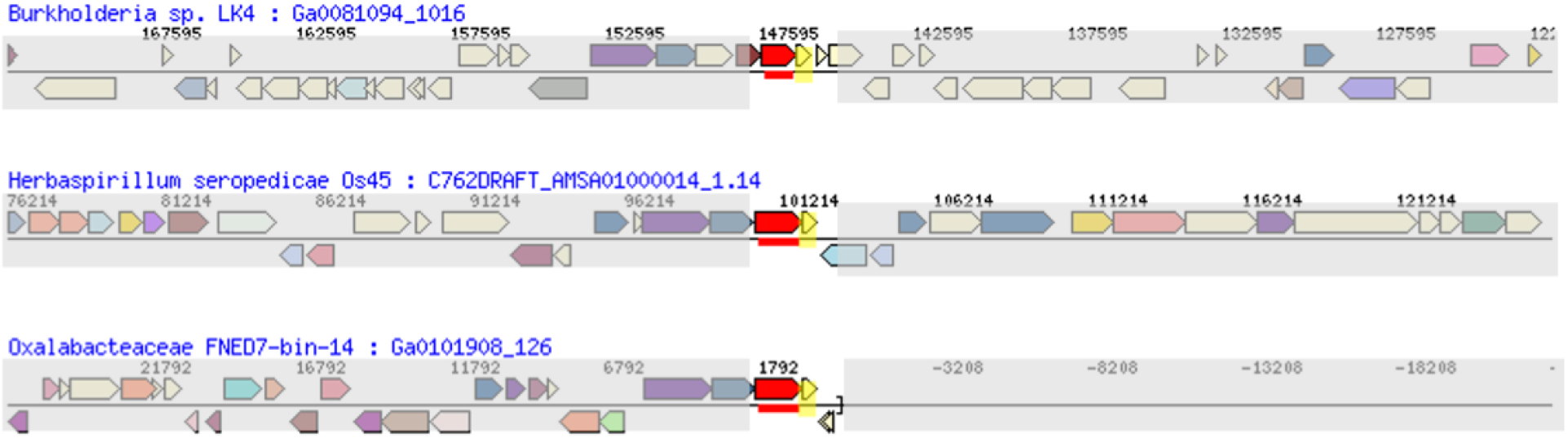
T6EC2 and T6IC2 display synteny across various genomes. Each row is a view of the noted genome (blue font). Red gene = T6EC2 and its homologs. Yellow = T6IC2 and its homologs.

**Supplementary Figure 13.**
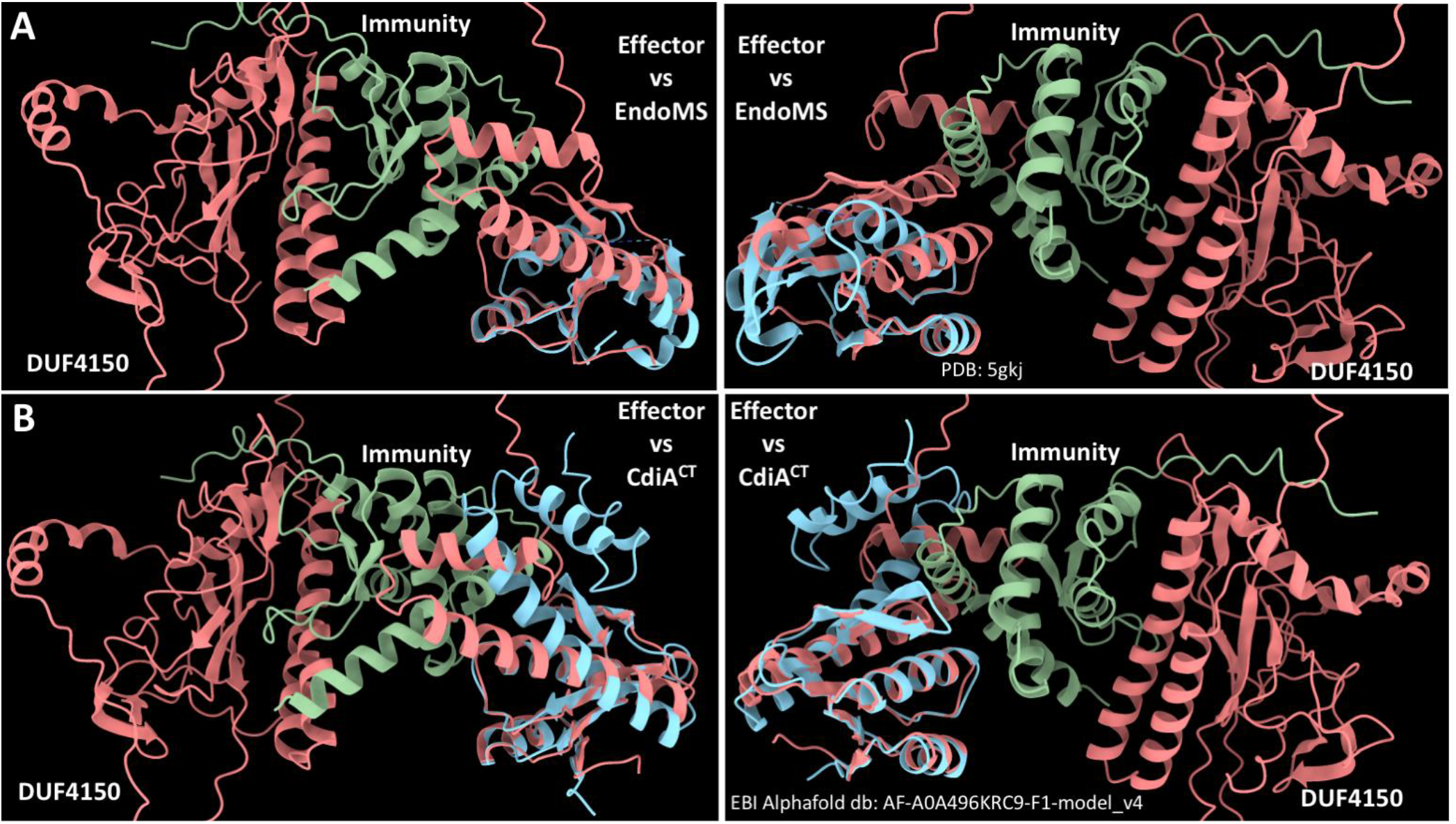
Structural search, but not sequence-based search, identified T6EC2 as a homolog with a C-terminal domain of EndoMS and CdiA. (A) Foldseek search of T6EC2 (pink) showed a partial match to EndoMS (PDB:5gkj). (B) Foldseek search of T6EC2 (pink) showed a match to the C-terminal domain of a CdiA contact-dependent inhibition protein (light blue; EBI Alphafold db: AF-A0A496KRC9-F1-model_v4). The protein structures were overlaid and visualized on ChimeraX.

**Supplementary Figure 14.**
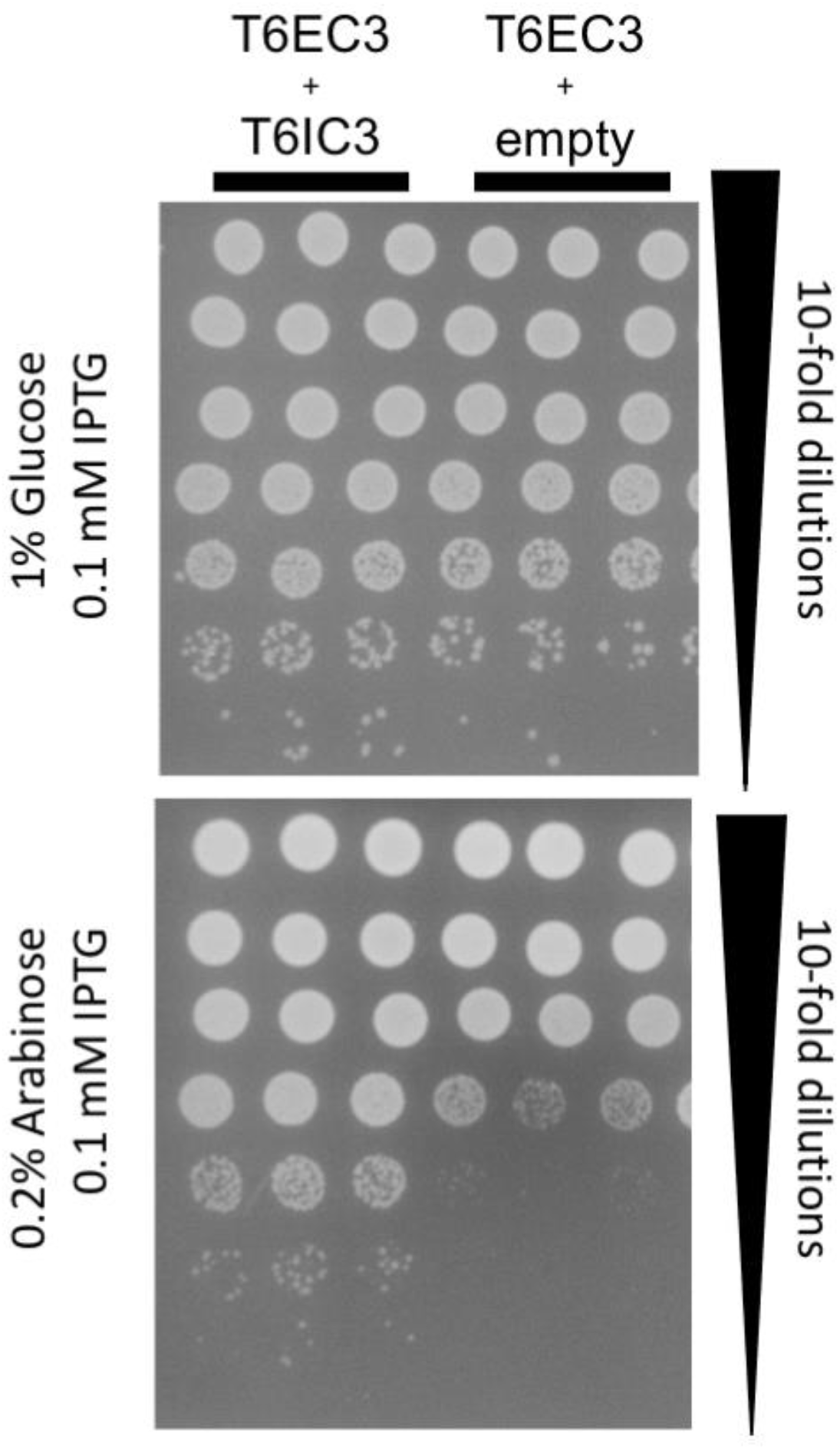
Heterologous expression of T6EC3 and T6IC3 genes in *E. coli* BL21. Drop assay of *E. coli* BL21 heterologously expressing T6EC3 in pBAD24 (arabinose induction) and T6IC3 in pET29b (IPTG induction), or T6EC3 in pBAD24 and an empty pET29b. T6EC3 is either uninduced (0.01 mM IPTG, 1% Glucose) or induced (0.01 mM IPTG, 0.2% Arabinose).

**Supplementary Figure 15.**
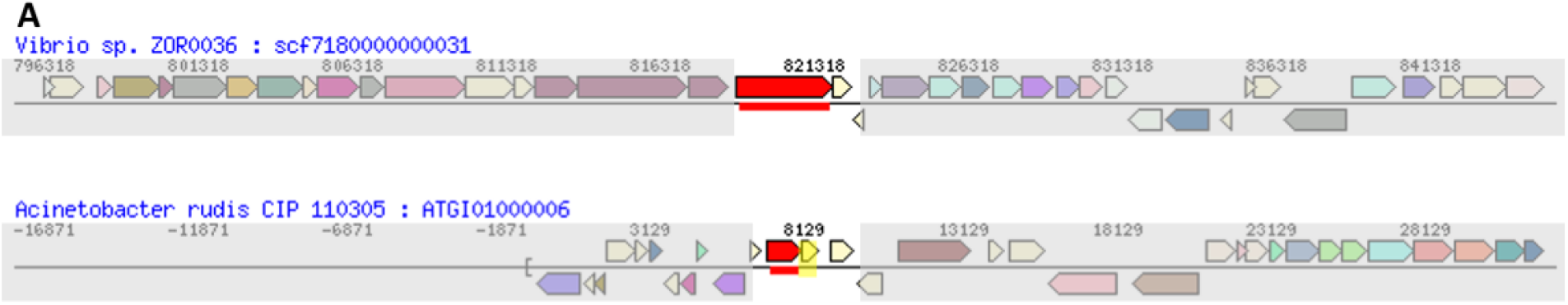
T6EC3 and T6IC3 display synteny across two genomes. Each row is a view of the noted genome (blue font). Red gene = T6EC3 and its homolog. Yellow = T6IC3 and its homolog. The top red gene is an “evolved” VgrG and also includes a peptidoglycan binding domain, in addition to the T6EC3 effector domain.

**Supplementary Figure 16.**
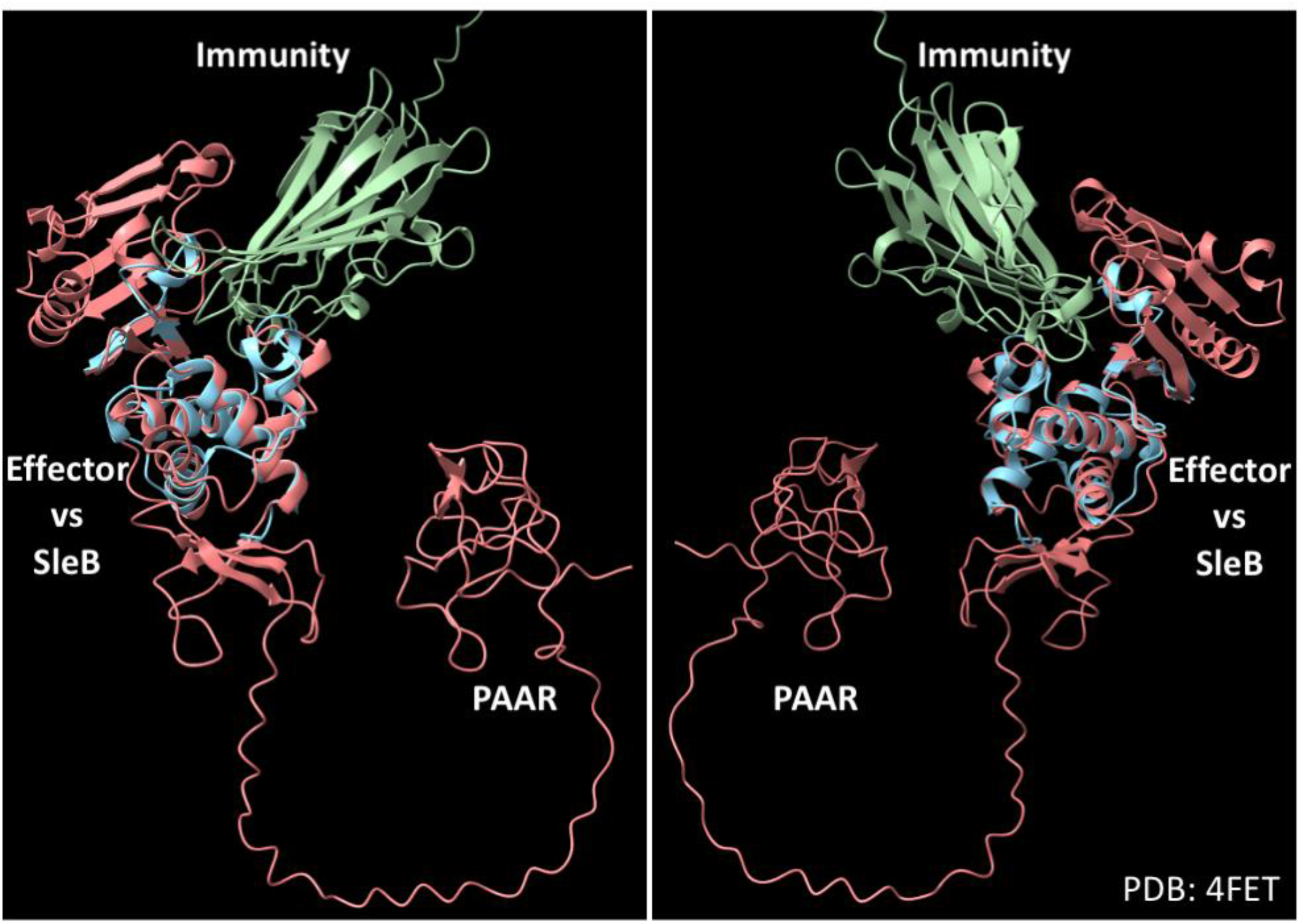
Structural search, but not sequence-based search, identified T6EC3 as a homolog of SleB. Foldseek search of T6EC3 (pink) showed a match to SleB (light blue; PDB accession: 4FET). The protein structures were overlaid and visualized on ChimeraX^61^.

**Supplementary Figure 17.**
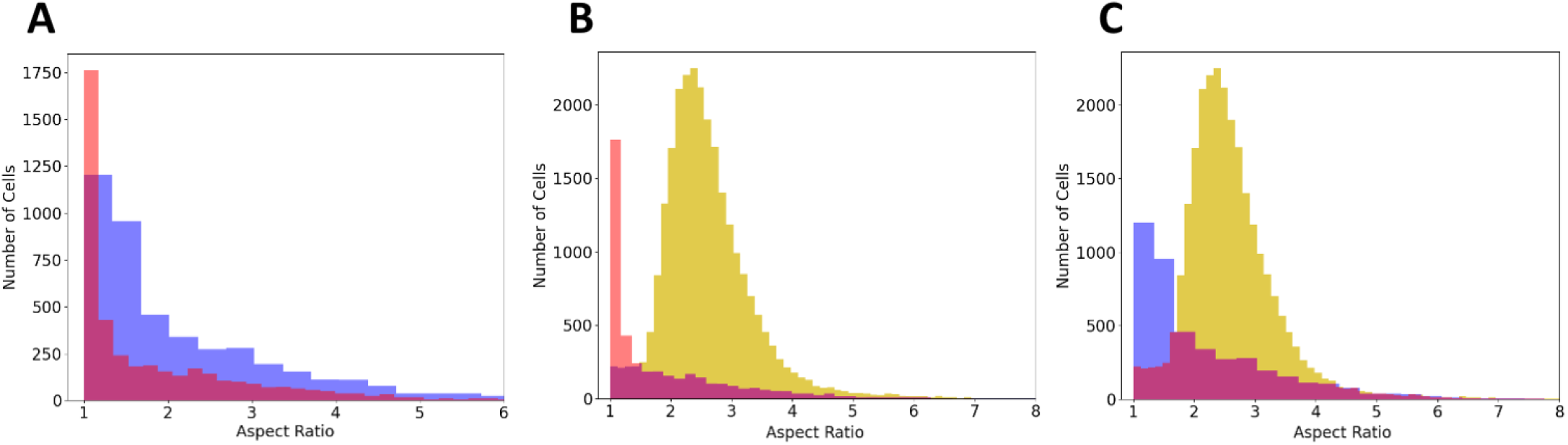
T6IC3 only inhibits rounding, but not to the level of wild type. (A) *E. coli* after 15 minutes of expression of T6EC3 with an empty vector (red) or T6EC3 and T6IC3 (blue). Differences between the two distributions were nonetheless significant(Mann-Whitney U statistic=11668854.0, pvalue=3.529e-104). (B) Expression of T6EC3 with an empty vector (red) versus *E. coli* with empty vectors (gold). (C) Expression of T6EC3 and T6IC3 (blue) versus *E. coli* with empty vectors (gold). Expression was induced with 0.01 mM IPTG and 0.2% Arabinose.

**Supplementary Figure 18.**
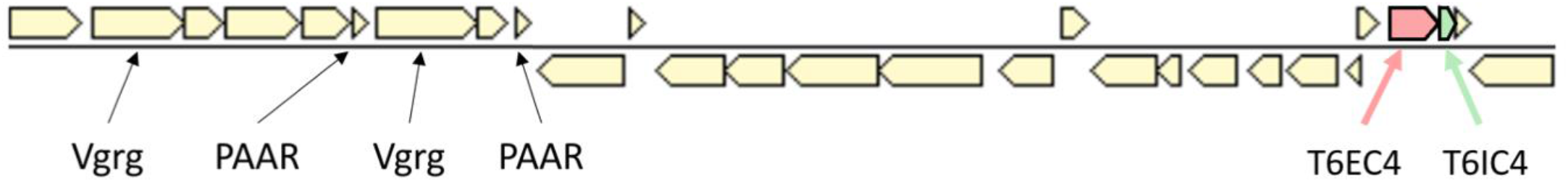
T6EC4 and T6IC4 are not part of a T6SS operon, although T6SS core genes are encoded nearby. The T6SS core genes are about 20,000 bp away from T6EC4 and T6IC4 (which are shown in pink and green, respectively).

**Supplementary Figure 19.**
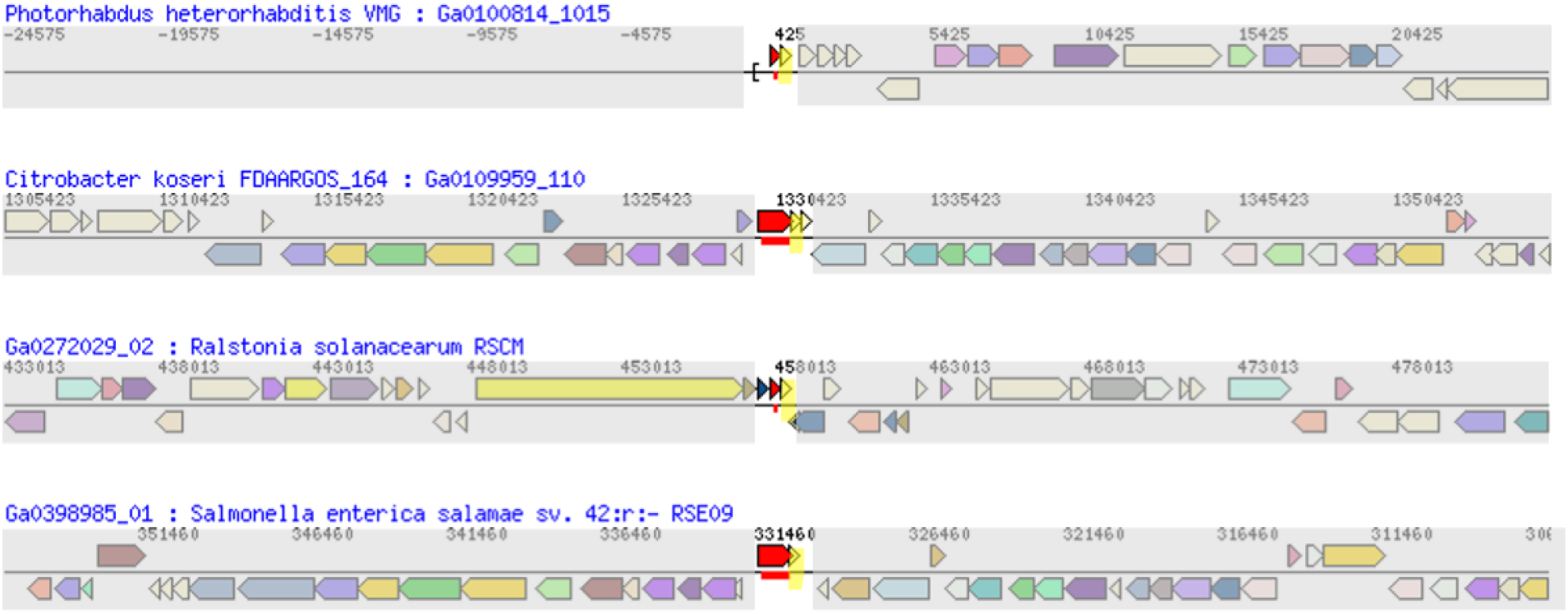
T6EC4 and T6IC4 display synteny across various genomes. Each row is a view of the noted genome (blue font). Red gene = T6EC4 and its homologs. Yellow = T6IC4 and its homologs.

**Supplementary Figure 20.**
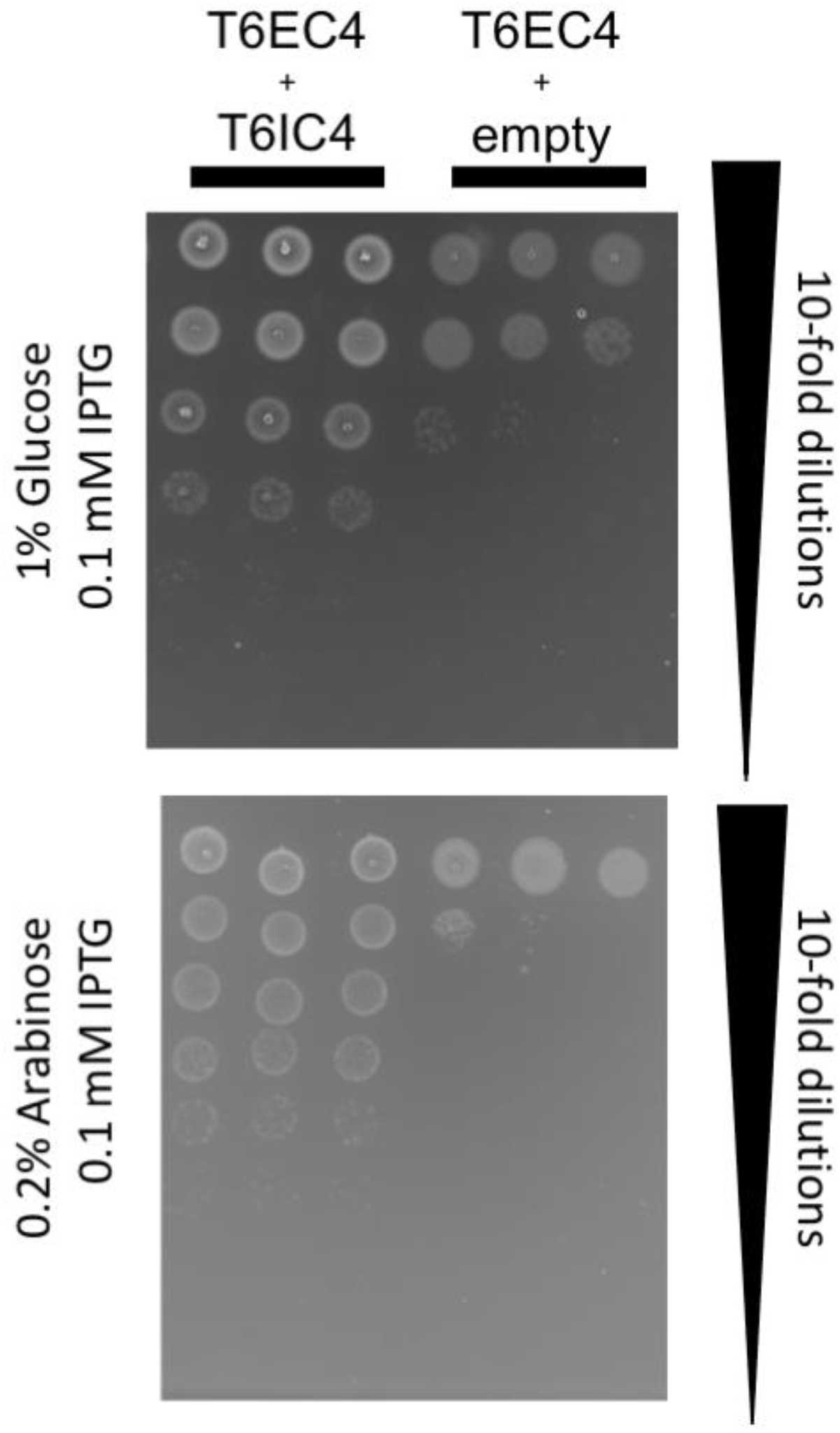
Heterologous expression of T6EC4 and T6IC4 genes in *E. coli* BL21. Drop assay of *E. coli* BL21 heterologously expressing T6EC4 in pBAD24 (arabinose induction) and T6IC4 in pET29b (IPTG induction), or T6EC4 in pBAD24 and an empty pET29b. T6EC4 is either uninduced (0.01 mM IPTG, 1% Glucose) or induced (0.01 mM IPTG, 0.2% Arabinose).

**Supplementary Figure 21.**
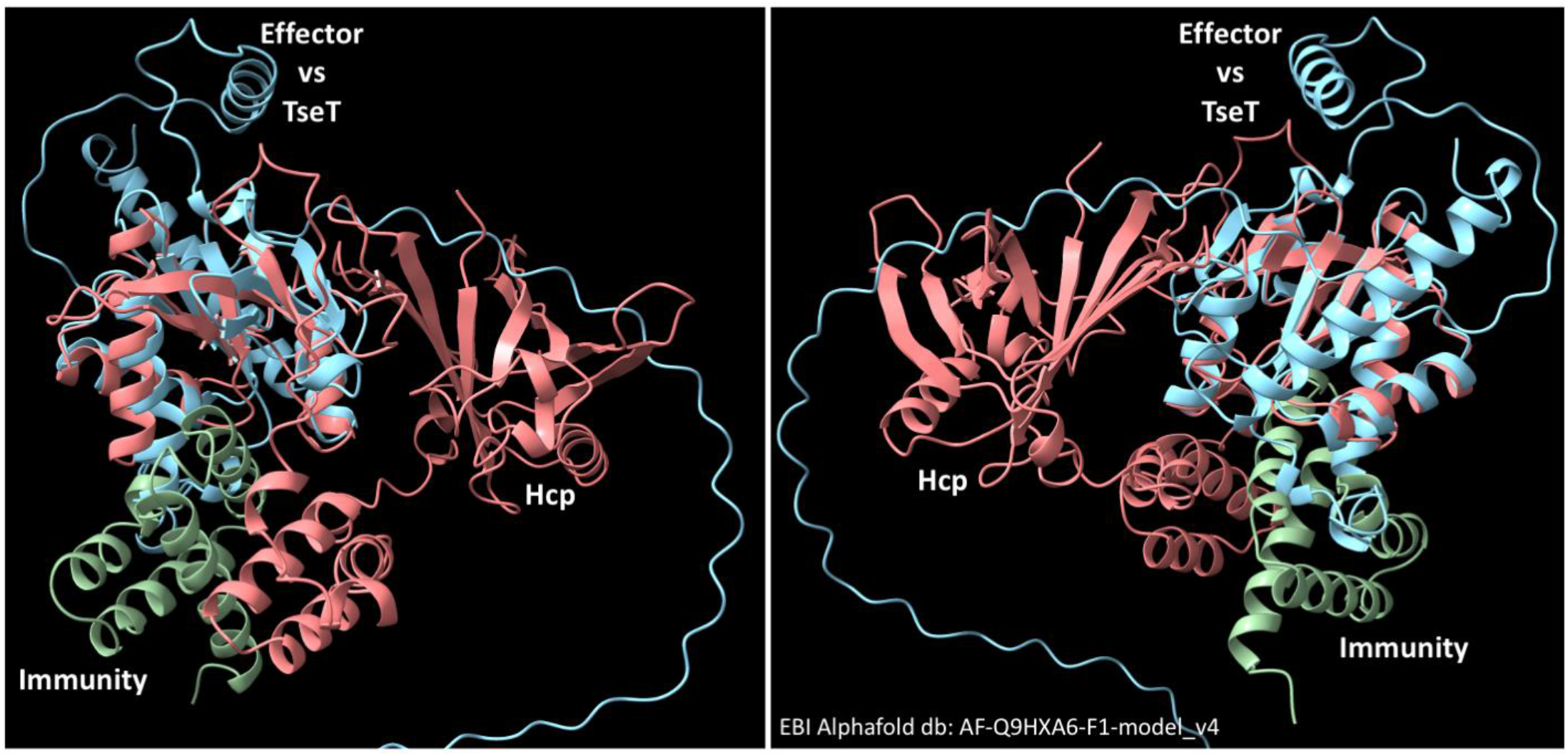
T6EC4 has structural similarity to T6E TseT from *Pseudomonas aeruginosa* PAO1. Foldseek search of T6EC4 (pink) showed a match to TseT (light blue; EBI Alphafold db: AF-Q9HXA6-F1-model_v4). The protein structures were overlaid and visualized on ChimeraX^61^.

